# Phage genome cleavage enables resuscitation from Cas13-induced bacterial dormancy

**DOI:** 10.1101/2022.07.05.498905

**Authors:** Madison C Williams, Alexandra E Reker, Shally R Margolis, Jingqiu Liao, Martin Wiedmann, Enrique R. Rojas, Alexander J Meeske

**Affiliations:** Department of Microbiology, University of Washington, Seattle, WA 98109, USA; Department of Civil and Environmental Engineering, Virginia Tech, Blacksburg, VA 24060, USA; Department of Food Science, Cornell University, Ithaca, NY, 10065, USA; Graduate Field of Microbiology, Cornell University, Ithaca, NY, 10065, USA; Department of Biology, New York University, New York, NY 10003, USA

**Keywords:** Cas13, restriction-modification, CRISPR, phage, dormancy

## Abstract

CRISPR-Cas systems provide their prokaryotic hosts with sequence-specific immunity to foreign genetic elements, including bacteriophages and plasmids. While most interfere with phage infection though cleavage of viral DNA, type VI CRISPR systems use the RNA-guided nuclease Cas13 to recognize mRNA targets. Upon engaging with target RNA, Cas13 cleaves both phage and host transcripts nonspecifically, leading to a state of cell dormancy that is incompatible with phage propagation. However, whether and how infected cells recover from dormancy is not clear. Here we show that type VI CRISPR systems frequently co-occur with DNA-cleaving restriction modification (RM) systems. Using genetics and microscopy, we show that Cas13 and RM systems synergize in anti-phage defense in the natural type VI CRISPR host *Listeria seeligeri*. Cleavage of the phage genome by RM removes the source of phage transcripts, enabling cells to recover from Cas13-induced cellular dormancy. We find that Cas13 and RM systems operating simultaneously eliminate phage DNA and neutralize infection more effectively than either defense alone. Thus, cells harboring both defense systems exhibit robust anti-phage immunity and survive infection. Our work therefore reveals that type VI CRISPR immunity is cell-autonomous and non-abortive, if paired with RM or similar DNA-targeting defenses. The ability of an abortive response to be resolved by the actions of another anti-phage defense has implications for the roles of diverse host-directed immune systems in bacteria.

## INTRODUCTION

In response to phage predation, bacteria have evolved elaborate repertoires of antiviral defenses that provide immunity through diverse mechanisms of action^1, 2^. During infection, activities of prokaryotic immune effectors and the nature of the invading virus both influence the fate of the infected cell. While some effectors elicit cell survival by direct neutralization of the virus (termed cell-autonomous immunity), others work via abortive infection, initiating a growth arrest or programmed cell death response in the infected cell, which promotes survival of uninfected kin. Many bacteria encode both abortive and non-abortive defenses that work simultaneously; how their combined activities affect the outcome of infection is not well understood.

CRISPR-Cas systems use RNA-guided nucleases to provide sequence-specific immunity against foreign genetic elements, including phages^3^, plasmids^4^, and transposons^5^. CRISPR loci contain arrays of 29-37 bp DNA repeats interspersed with unique spacer sequences of foreign origin^6, 7^, as well as *cas* genes encoding effector nucleases^8, 9^. During CRISPR immunity, small fragments of foreign nucleic acids are captured and integrated into the CRISPR array, where they serve as immunological memories for future infection^3^. The CRISPR array is transcribed and processed into small crRNAs, which associate with Cas nucleases and arm them for surveillance and cleavage of complementary nucleic acids^10–12^. Bioinformatic analyses have uncovered diverse CRISPR-Cas systems, which are classified into 6 types based on their *cas* gene families and interference mechanisms^9^. While most CRISPR-Cas systems act by cleavage of phage DNA, the type VI CRISPR nuclease Cas13 exclusively recognizes and cleaves RNA^13^. We previously discovered that, upon engagement with targeted phage mRNA, Cas13 is activated as a non-specific RNase and degrades both phage and host RNA in its natural host bacterium *Listeria seeligeri*^14^. Under these circumstances, cleavage of host RNAs results in a state of cellular dormancy incompatible with either continued cell growth or progression of the phage lytic cycle. Type VI CRISPR-Cas systems can provide robust anti-viral immunity to a wide range of viruses through this abortive infection-like mechanism^13–17^.

In addition to CRISPR immunity, roughly 74% of bacterial genomes encode restriction- modification (RM) systems, which use restriction endonucleases to recognize and cleave 4-8 bp motifs on phage DNA^18–20^. Adenine or cytosine methylases encoded by RM loci modify these sites on the bacterial genome to protect it from autoimmune cleavage. RM systems are classified into 4 types: type I RM proteins form multiprotein complexes comprised of nuclease (R), methylase (M), and specificity (S) subunits.

Specificity subunits are DNA-binding proteins that determine the RM recognition sequence, and can alternately associate with other subunits as an M2S trimer to perform site methylation, or as an M2R2S pentamer to perform cleavage of unmethylated sites. The R subunit translocates a variable distance away from the unmethylated target site and generates a double stranded break. In contrast, type II RM nucleases and methylases act separately on DNA, and cleavage is performed directly at the recognition site. Type III RM systems combine elements of types I and II, and are composed of multisubunit complexes that cleave at a fixed distance from the target site. Type IV RM systems do not encode a methylase and instead cleave modified DNA sites. Cleavage of phage DNA by RM systems is generally thought to enable survival of the infected cell, making RM a cell-autonomous immune mechanism. Finally, methylases that modify and protect host DNA can sometimes methylate recognition sites on phage DNA, rendering them refractory to nucleolytic cleavage^20^. Phage DNA methylation is a common mechanism by which RM immunity is subverted.

Although many bacteria that encode type VI CRISPR systems likely also carry one or more RM systems, there has been no investigation into how the activities of these systems influence one another. Two reports have demonstrated that DNA-targeting CRISPR systems and RM can exert anti-phage immunity at the same time^21, 22^. Previous investigations of type VI CRISPR immunity have been performed with methylated or otherwise restriction-resistant phages^13–17^. Here we find that RM systems frequently co- occur with type VI-A CRISPR systems in strains of *L. seeligeri* and other *Listeria*, and the two defenses operate simultaneously during phage infection. Our experimental data indicate that RM-mediated cleavage of phage genomes enables the survival and growth of infected cells, despite activation of nonspecific RNA cleavage by Cas13. Finally, we observed that the clearance of phage DNA, neutralization of infection, and cell growth after infection are all more effective in cells equipped with both RM and Cas13 than strains harboring either defense alone. While type VI CRISPR systems can provide effective immunity through abortive infection, our work indicates that they also enhance the cell-autonomous immunity elicited by DNA-targeting defenses.

## RESULTS

### Type VI CRISPR systems frequently co-occur with Restriction Modification systems

Phylogenetic analysis of Cas13 homologs suggests that *Listeria* is a major clade harboring type VI-A CRISPR systems^23^. We recently established the natural type VI-A CRISPR host *Listeria seeligeri* as a genetically tractable model for studying Cas13a- based immunity^14, 17, 24^. To better understand the context in which type VI CRISPR systems function, we sequenced the genomes of 62 diverse isolates of *L. seeligeri*, and evaluated their anti-phage defense system content. We searched for components of CRISPR, restriction-modification (RM), BREX, Abi, DISARM, CBASS, Pycsar, and Deity systems^25^. Of these defenses, CRISPR and RM systems were by far the most well- represented in *L. seeligeri* genomes (Fig S1A). We identified four CRISPR types in *L. seeligeri*: DNA-targeting type I-B (in 69% of genomes), type II-A (50%), type II-C (6%) loci as well as RNA-targeting type VI-A loci (in 29% of genomes). We observed these CRISPR loci in diverse combinations (Fig. 1B), while some strains possessed only one CRISPR type, most harbored multiple types, and no two types co-occurred with each other exclusively. While most DNA-targeting CRISPR loci were found in 1-2 hotspots across different genomes, type VI loci exhibited larger variation in location (Fig S1B). Our genomic analysis also identified 4 different types of RM systems across the genomes (Fig S1C). 90% of the strains had one or more RM systems, including type I (62%), type II (45%), type III (6%), and type IV (42%). Like the CRISPR loci, RM loci were distributed in different locations across the genomes and found in a variety of combinations (Fig S1D). None of the RM loci were closely linked to CRISPR loci. Collectively, these observations suggest that anti-phage defense loci are rapidly evolving in *L. seeligeri*, possibly via horizontal gene transfer.

**Figure 1.**
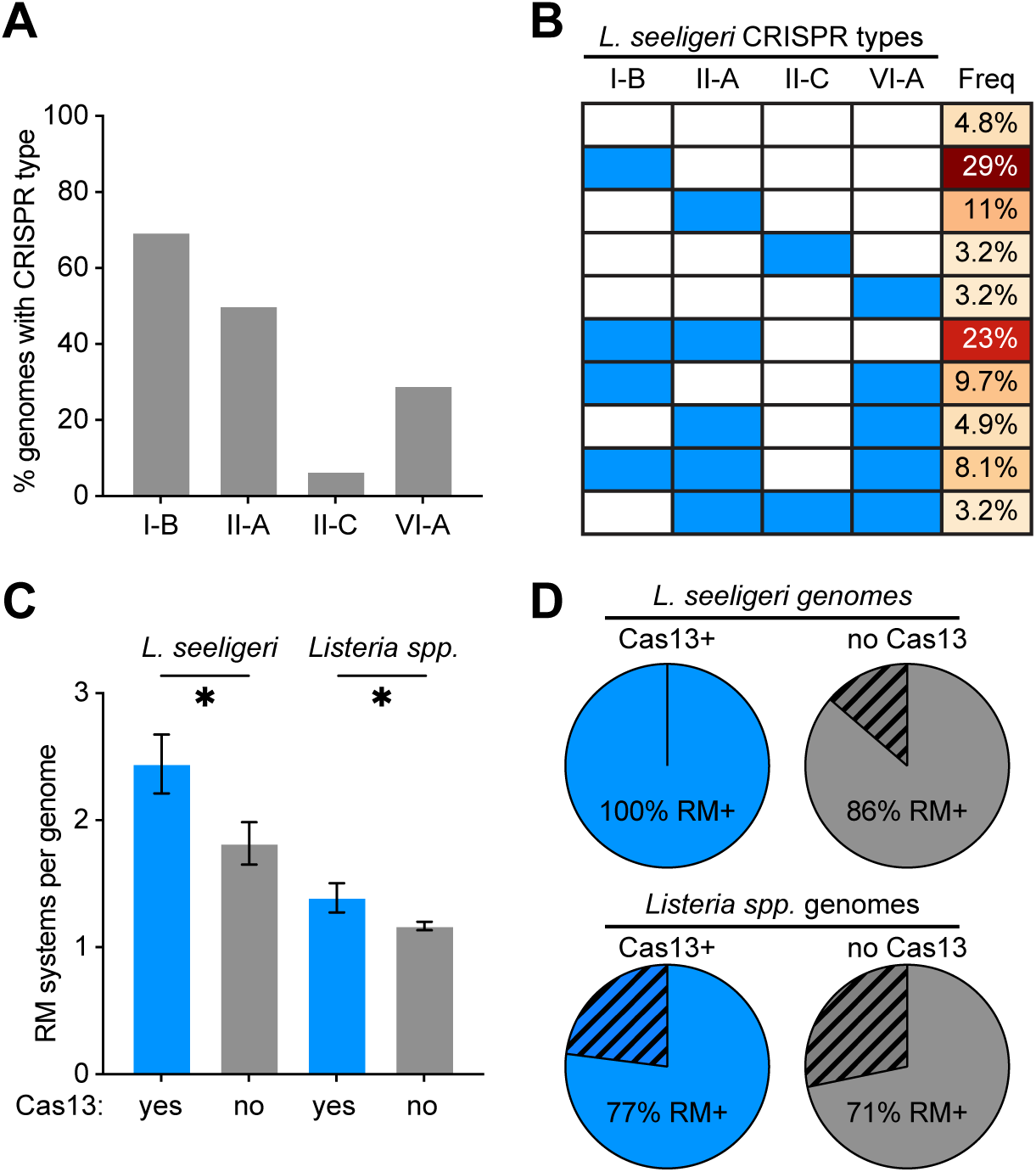
Type VI-A CRISPR systems frequently co-occur with RM systems in *Listeria* genomes. **(A)** Percentage of 62 *L. seeligeri* genomes harboring the indicated CRISPR type. **(B)** Combinations of CRISPR loci observed in 62 *L. seeligeri* genomes. Filled blue rectangles indicate the presence of the indicated CRISPR type. **(C)** Average tally of RM systems per genome and **(D)** percentage of genomes containing any RM loci in *Listeria* strains with or without Cas13. 62 *L. seeligeri* genomes and 943 genomes from a broader dataset of *Listeria spp*. were analyzed. Error bars denote standard error of the mean. Asterisks denote statistical significance (P<0.05) using Student’s *t-*test (two-tailed).

We next investigated whether any defense systems were closely associated with type VI CRISPR systems. We tallied the number of RM systems in each genome, and discovered a statistically significant enrichment in average number of RM loci within genomes also harboring type VI CRISPR loci (Fig 1C). In contrast, we detected no significant difference in RM content for strains with type I-B or type II-A CRISPR loci (type II-C loci were too sparse for a meaningful comparison) (Fig S1E). In addition to a higher number of RM loci, we found that strains with type VI CRISPR were also more likely to co-occur with RM systems in general, with 100% of type VI+ strains harboring at least one RM system (Fig 1D). Next, we expanded our analysis to include 943 *Listeria* genomes available in the NCBI whole genome sequencing database. Among these genomes, we detected a smaller but still statistically significant enrichment in RM co-occurrence and count in strains containing a type VI system. These bioinformatic analyses suggest that most *Listeria* type VI CRISPR loci could perform anti-phage defense in concert with one or more DNA-targeting RM systems.

### *L. seeligeri* LS1 has two functional type I RM systems

To investigate whether the RM systems we identified function in anti-phage immunity, we performed phage-challenge experiments in our most well-characterized strain, L. seeligeri LS1. The LS1 genome encodes two type I RM systems, each containing a methylase, specificity subunit, and nuclease. One of the two loci is also associated with a type IV RM nuclease (Fig S2). We examined the expression of these two genomic regions in our previously obtained RNA-seq dataset^14^, and found that both RM systems are transcribed at high levels during exponential growth (Fig S2). We deleted each RM locus individually, and also generated a double deletion lacking both loci. We performed efficiency-of-plaquing assays in which we challenged each of these mutants and wild-type LS1 with five different listeriaphages from our collection: A118, ϕEGDe, U153, ϕLS46, and ϕLS59 (Fig 2A). Importantly, LS1 has type I-B and type VI-A CRISPR-cas systems, but does not carry any spacers matching any of the phages tested, therefore CRISPR immunity was not activated during these experiments. Each RM system reduced plaquing efficiency for all five phages. Compared to the double RM deletion, the individual RM systems conferred between a 2-fold and 500-fold degree of protection, and both systems together provided even more immunity. To understand the basis of the variation we observed in RM-based immunity against different phages, we identified the DNA motifs recognized by each of the two RM systems. By performing PacBio sequencing of the LS1 genome in strains possessing each RM system individually, we identified 362 methylated adenines (m6A) in a strain with RM1 alone, and 620 methylation sites with RM2 alone. We aligned the surrounding sequences to generate consensus recognition motifs (Fig 2B, Supplementary Files 1-2). Consistent with other type I RM sites^19^, these motifs are each composed of two 4-5 basepair motifs separated by 7-8 random nucleotides. When we tallied the number of sites present in the genomes of the phages used for our efficiency-of-plaquing assays, we observed a direct correlation between the number of RM sites and degree of anti-phage defense (Fig 2C-D). Collectively, these data indicate that both type I RM systems of *L. seeligeri* LS1 use target site recognition to perform anti-phage defense against a variety of phages.

**Figure 2.**
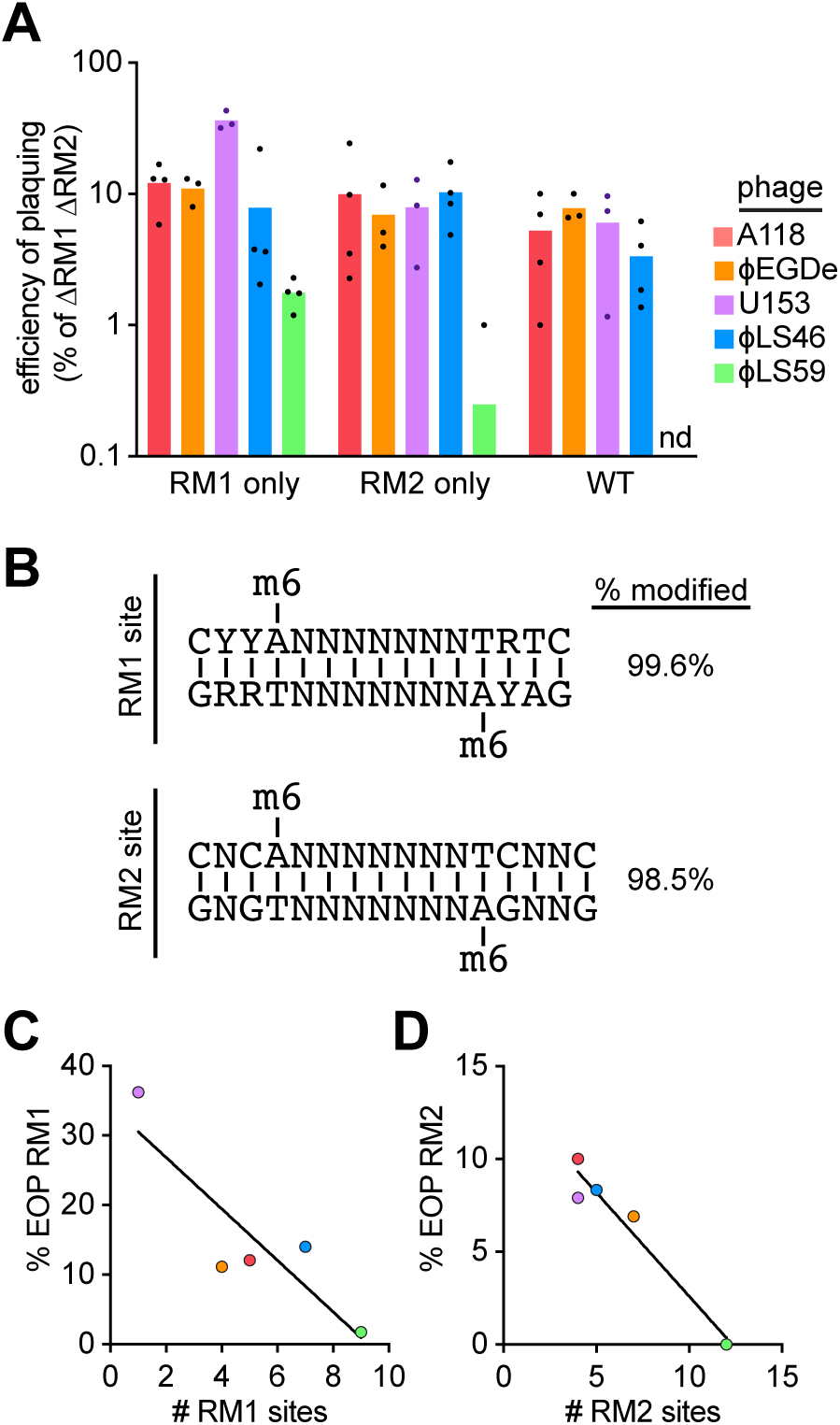
*L. seeligeri* LS1 encodes two functional type I RM systems. **(A)** Efficiency of plaquing assay testing ability of five different phages to infect LS1 strains with 0, 1, or 2 RM systems. Efficiency of plaquing values for each phage are reported as a percentage of the number of plaques formed on LS1 lacking both RM systems (ΔRM1 ΔRM2). ND = not detected. **(B)** Type I RM recognition motifs for LS1 RM1 and RM2 systems identified by PacBio sequencing. 6-methyl adenine modifications were detected at the indicated consensus motifs in genomic DNA purified from LS1 lacking RM2 (to identify the RM1 site) or lacking RM1 (to identify the RM2 site). The percentage of methylated genomic motifs matching the consensus are indicated on the right. Y=pyrimidine, R=purine. **(C)** Negative correlation between the number of RM1 recognition sites on phage genomes and the mean efficiency of plaquing (EOP) for that phage on bacteria harboring RM1. Each phage is color-coded as in (A). **(D)** Same as (C), but for RM2.

### RM systems allow survival of phage infection during type VI CRISPR immunity

During type VI CRISPR immunity, Cas13 recognition of crRNA-complementary phage mRNA triggers nonspecific *trans*-cleavage of both phage and host RNAs, leading to dormancy of the infected cell^13, 14^. As the RNase activity of Cas13 does not eliminate the phage DNA, which is the source of target RNA, infected cells remain dormant despite Cas13 activation. We therefore hypothesized that RM cleavage of phage genomes might allow infected cells to escape Cas13-induced cell dormancy. On the other hand, triggering of Cas13 activity might lead to an irreversible commitment to cell dormancy, regardless of phage genome elimination by RM. To explore this idea, we grew cultures of wild-type LS1 and the double RM deletion mutant in the presence or absence of a type VI spacer targeting the restriction-sensitive phage ϕLS59, and measured the number of viable CFU before and at 0.5, 4, and 24 hours post-infection (Fig 3A). The lysis time of phiLS59 is 2 hours, therefore the CFU measurement at 0.5h reflects the fates of the initially infected cells. We used a high MOI (10) to ensure that most cells were infected. We performed this experiment with two different spacers, one (spcE) targeting an early-transcribed ϕLS59 lytic gene, and one (spcL) targeting a late- transcribed gene, each of which provide robust immunity against ϕLS59 (Fig S3). Like our previous observations with other phage^14^, we observed an initial ∼50-fold reduction in CFU after infecting cells lacking RM immunity, regardless of the presence of a targeting spacer, indicating that infected cells are mostly unable to form a viable colony. After 24 hours, while ΔRM cultures with a non-targeting spacer lysed to near completion, cultures with a targeting spacer recovered and proliferated. In contrast, all three of the wild-type strains containing RM systems (with or without spcE and spcL) maintained viability after infection, and all continued to proliferate over the time course.

**Figure 3.**
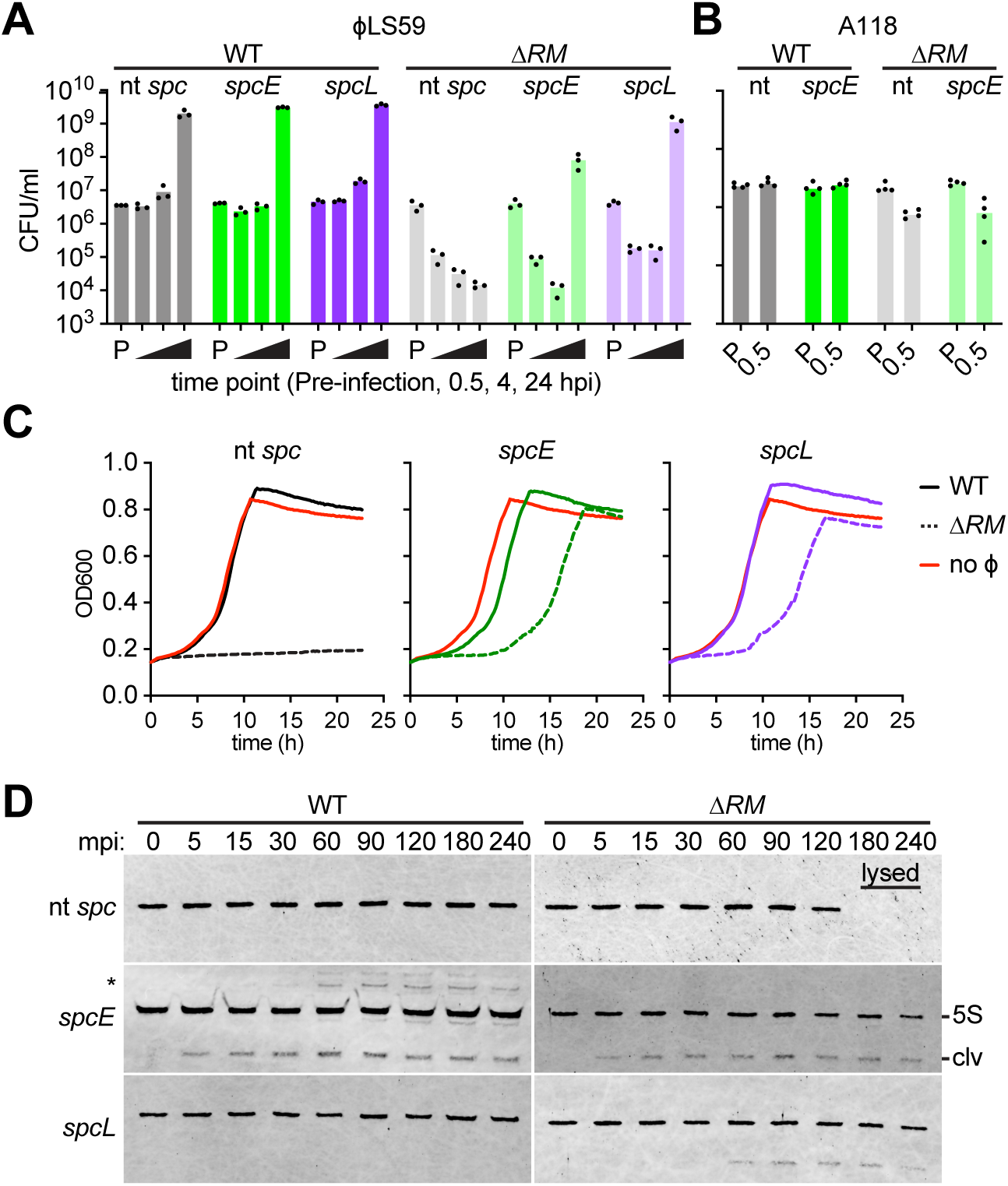
RM systems enable survival of phage infection during type VI CRISPR immunity. **(A)** Viable colony-forming-unit (CFU) count during ϕLS59 infection of *L. seeligeri* LS1. Cells of the indicated genotype were infected ϕLS59 at MOI 10. Viable CFU were enumerated prior to infection (P), as well as 0.5, 4, and 24 hours post- infection. NT = non-targeting spacer, spcE = spacer targeting early lytic gene, spcL = spacer targeting late lytic gene. **(B)** CFU count during A118 infection of LS1 at MOI 10. **(C)** Growth curves during ϕLS59 infection of LS1. Strains of the indicated genotype were infected at MOI=10, OD600=0.01. OD600 was monitored for 24 hours after infection. Solid lines indicate wild-type, RM^+^ strains, dashed lines indicate ΔRM strains. Uninfected controls are shown in red. The mean of 3 biological replicates is plotted. **(D)** Northern blot analysis of 5S rRNA cleavage during type VI CRISPR immunity. Strains of the indicated genotype were infected with ϕLS59 at MOI 10, then RNA was extracted and probed for 5S rRNA. Cas13-dependent cleavage products (clv) and putative rRNA precursors (*) are shown. Representative of two biological replicates.

We observed a similar survival phenotype during infection by a different phage, A118, in the presence of both RM and an early-targeting spacer (Fig 3B). These results suggest that cells endowed with both RM and type VI CRISPR immunity largely survive infection.

We considered that cells equipped with both RM and type VI CRISPR immunity could survive phage infection by two mechanisms: (i) RM nucleases could cleave the phage genome before transcription of the phage lytic genes, therefore Cas13 activity would never be triggered and the cells would not enter dormancy, or (ii) phage lytic gene transcription, Cas13 activation, and cell dormancy all precede phage genome cleavage by RM nucleases, but elimination of the phage genome allows the cell to be resuscitated. To distinguish between these possibilities, we determined whether Cas13 is activated in the presence of RM immunity during phage infection. First, we analyzed cell growth rates during ϕLS59 infection of cells containing RM, type VI targeting spacers, or both (Fig 3C). We infected cells at MOI 10 and tracked culture growth (OD600) over a period of 24 hours. Consistent with the CFU survival data in Fig 3A, cells lacking both RM and CRISPR immunity were lysed by the phage, and did not recover within the growth period. In contrast, cells harboring RM but with a non- targeting spacer grew at rates indistinguishable from uninfected cells. The strains lacking RM but armed with a type VI spacer initially exhibited severe growth defects after infection, but later recovered. Our CFU measurements indicated that most infected cells in this population were unable to form colonies, therefore uninfected cells were likely responsible for the eventual regrowth of the culture. Finally, while the strain containing both RM and spcL exhibited unperturbed growth, we observed a small but reproducible lag in growth after infection of the strain containing both RM and spcE. Consistent with these measurements, we noted a similar trend in our CFU measurements (Fig 3A) at 4 hours post-infection: the wild-type RM^+^ strain with either a non-targeting spacer or spcL proliferated more at this time point than the strain armed with spcE. These results suggest that cells equipped with spcE trigger Cas13 immunity and cell dormancy before RM systems can clear infecting phage genomes.

To directly test whether Cas13 is activated during ϕLS59 infection in the presence of RM systems, we performed northern blots that enabled us to monitor Cas13-mediated RNA cleavage over time in vivo. We blotted for the L. seeligeri 5S rRNA, an extremely abundant RNA that we previously demonstrated is a substrate of the trans-RNase activity of Cas13 that is stimulated upon target RNA recognition^24^. In the absence of RM or CRISPR immunity, we observed a single prominent 5S rRNA band throughout the course of infection. For strains lacking RM but carrying a type VI targeting spacer, we observed the formation of lower molecular weight cleavage products after phage infection. The onset of cleavage depended on the timing of target expression: for spcE, cleavage was evident at 5 minutes post-infection, for spcL, cleavage was delayed until 60 minutes post-infection. In contrast, in the wild-type RM^+^ strains with either a non- targeting spacer or spcL, 5S rRNA remained intact for at least 4 hours after infection, mirroring the growth curve data in Fig 3C and suggesting Cas13 was not activated in these strains. Finally, the RM^+^ strain harboring spcE exhibited rapid-onset cleavage of 5S rRNA after infection, indicating that Cas13 is activated prior to RM-mediated clearance of the phage genome. Importantly, the extent of cleavage was similar for spcE^+^ cells, regardless of the presence of RM, suggesting a similar degree of Cas13 activity in both strains. We also noted the formation of higher molecular weight bands during later time points in the RM^+^ spcE^+^ cells. We hypothesize that these bands represent the precursor rRNAs that are processed to generate mature 5S rRNA^26^. The formation of these intermediates may be a consequence of resuscitation from Cas13- induced dormancy. Collectively, these results suggest that type VI CRISPR immunity can either precede or follow RM cleavage of the phage genome, depending on the timing of target transcript expression. Furthermore, our observation of cell survival despite Cas13 activation during infection of the RM^+^ spcE^+^ strain suggests that RM- mediated cleavage of the phage genome allows cells to exit Cas13-induced dormancy.

### RM cleavage of phage genomes enables recovery from Cas13-induced dormancy

To directly test whether RM systems allow exit from Cas13-mediated cell dormancy, we monitored the growth of single cells after infection via microscopy. We used ϕLS59 to infect wild-type and ΔRM strains with or without a targeting spacer at an MOI of 10. Then, we loaded infected cells into microfluidic chambers where they were trapped for imaging. After a 10 minute adsorption period, we washed out unbound phage with growth medium, which was supplied continuously throughout the experiment. To ensure that the majority of cells were stably infected with under these conditions, we labeled ϕLS59 by soaking in the nucleic acid stain SYTOX Green (Fig 4A). Because the stain is membrane-impermeable, it is only delivered to cells via genome injection, and is not taken up by uninfected cells^27^. During the early stages of infection with labeled phage, fluorescent phage particles were visible at the cell periphery. Within 20 minutes, infected cells exhibited bright cytosolic fluorescence as a consequence of injection of the labeled phage genome. When we imaged cells infected at MOI 10, virtually every cell contained fluorescent label (Fig 4B). Next, we monitored cell growth for 5 hours after infection. We performed these experiments with unlabeled phage to avoid the effects of phototoxicity or any artifactual impact from the nucleic acid stain. After phage infection, we observed 4 distinct phenotypes (Fig 4C-D). Cells lacking RM and CRISPR grew unperturbed, but almost always lysed within 3 hours after infection. In contrast, ΔRM cells containing spcE exhibited rapid growth arrest. While nearly every cell ceased growth, 23% eventually lysed during the timelapse, suggesting that Cas13-induced dormancy can be lethal under some circumstances. In the presence of RM immunity alone, nearly all cells grew and divided normally after infection, and only rarely lysed or entered stasis. Finally, RM^+^ cells armed with spcE exhibited a range of phenotypes in our experiments. While 36% of the cells grew without delay, 32% underwent transient growth arrest but then resumed growth. Finally, 27% remained in a state of arrest throughout the imaging period. We noted two forms of recovery from dormancy that would each result in the formation of a viable colony: (i) infected cells that underwent delayed growth, then began elongating and dividing again, (Fig 4C, 4^th^ column) and (ii) infected cells that divided into two daughter cells, one of which continued growth, while the other stayed in arrest (Fig 4C, 5^th^ column). These microscopy data directly demonstrate that cells can use RM immunity to survive the collateral effects of Cas13 RNase activities during type VI CRISPR immunity.

**Figure 4.**
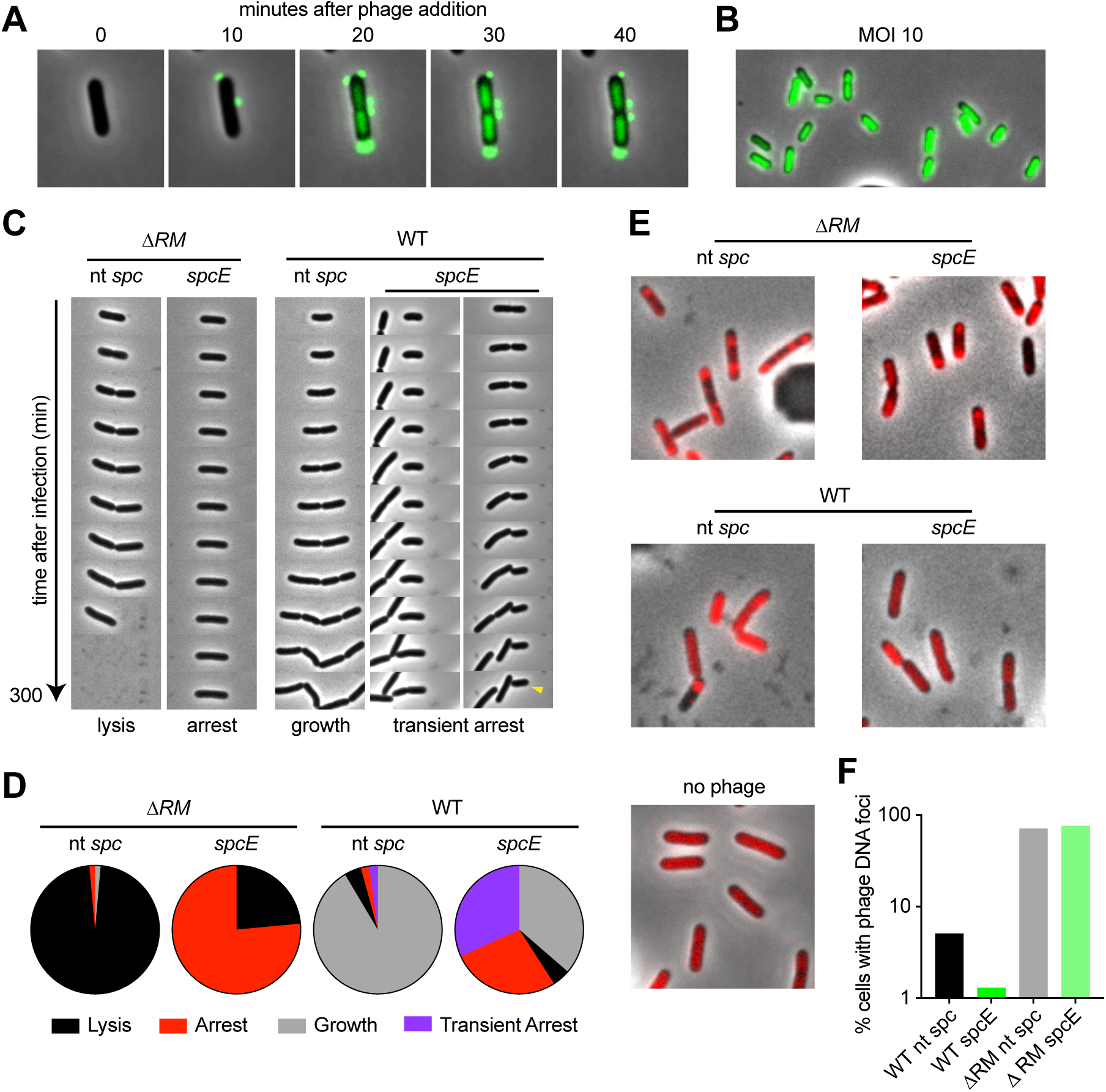
RM cleavage of phage genomes enables recovery from Cas13-induced cell dormancy. **(A)** Timelapse montage of LS1 cells infected with Sytox Green-labeled ϕLS59 in a microfluidic chamber. Overlay of SYTOX Green signal and phase contrast images. **(B)** Field of cells infected with SYTOX Green-labeled ϕLS59 at MOI 10 and loaded into microfluidic chamber. **(C)** Timelapse montage examples showing cells of the indicated genotype during growth in a microfluidic chamber after ϕLS59 infection. The fate of each cell is indicated below. For WT+*spcE*, two example phenotypes are shown. Left, a cell exhibiting delayed growth. Right, a cell that divides into a growing daughter and a dormant daughter (indicated with yellow caret). **(D)** Percentage of cells exhibiting the indicated fate for each genotype. 500 cells were analyzed for each genotype. **(E)** Representative images from FROS experiments tracking ϕLS59 DNA during infection. LS1 cells harboring tetR-mCherry were infected with ϕLS59 tetOx60 and imaged 40 minutes post-infection. **(F)** Quantitation of data in E; percentage of cells of each genotype with detectable phage DNA foci. 500 cells were analyzed for each genotype.

We hypothesized that RM systems enable escape from Cas13-mediated dormancy by eliminating phage DNA. Therefore, we wanted to test whether clearance of phage DNA correlated with cellular recovery during infection. We were unable to use SYTOX labeling to specifically label phage DNA, because after phage genome injection the nucleic acid stain labels both the phage and bacterial genomes. To track the presence of ϕLS59 DNA after infection, we developed a Fluorescent Reporter-Operator System (FROS)^28^ using an mCherry-labeled allele of the DNA binding protein TetR and its 19 bp binding motif *tetO*. Then, we generated a ϕLS59 mutant harboring an array of 60 *tetO* sites inserted downstream of the late lytic genes. We confirmed that this mutant phage retained sensitivity to both RM and Cas13 immunity (Fig S4). In the absence of phage infection, we observed a uniformly distributed cytosolic fluorescent signal in cells carrying a plasmid-encoded tetR-mCherry (Fig 4E). Next, we infected ΔRM cells without a targeting spacer using ϕLS59 tetOx60. After infection, the tetR-mCherry signal in most cells coalesced into 1-2 foci, usually positioned towards the cell poles (Fig 4E-F). When we infected a ΔRM strain equipped with spcE and tetR-mCherry, we observed a similar degree of phage genome foci formation, suggesting that Cas13 immunity is insufficient to eliminate the phage genome. In contrast, when we infected RM^+^ cells with ϕLS59 tetOx60, most cells did not exhibit phage genome foci, instead displaying diffuse cytosolic fluorescence similar to our observations of uninfected cells. Notably, we did observe instances of phage genome foci in 5% of RM^+^ cells lacking Cas13 immunity. Phage genome foci were even rarer (1% of cells) in RM^+^ spcE cells, suggesting that although Cas13 does not directly interfere with phage DNA, Cas13 activities enhance the clearance of phage DNA during RM immunity. Collectively, these observations indicate that Cas13 activity does not lead to an irreversible stasis in cell growth, but RM clearance of phage genomes allows survival and recovery from Cas13-induced dormancy during type VI CRISPR immunity.

### RM and type VI CRISPR systems synergize in anti-phage immunity

While the results described above demonstrate that RM and Cas13 immunity can work simultaneously, they do not explain what advantage type VI CRISPR immunity might confer to bacteria already armed with RM. On the contrary, Cas13-induced dormancy actually elicited a growth disadvantage during infection with ϕLS59 (Fig 3C). To investigate this, we measured the number of productive phage infections using a highly sensitive center of infection assay (Fig 5A). We infected wild-type and ΔRM cells with or without *spcE* with ϕLS59 at MOI 10, thoroughly washed them to remove remaining phage particles, then plated the infected cells (centers of infection) on a lawn of phage- sensitive ΔRM cells lacking CRISPR. In this assay, infected cells that successfully produce even a single phage particle will give rise to a detectable plaque. Despite our observation that ϕLS59 does not form plaques on wild-type LS1 (Fig 2A), we observed that ϕLS59 productively infects ∼1% of cells containing either RM or Cas13+*spcE* alone. In contrast, we were unable to detect any successful infections of cells harboring both RM and type VI CRISPR immunity. Thus, while Cas13 immunity elicits a growth defect during anti-phage immunity, it synergizes with RM in phage neutralization.

**Figure 5.**
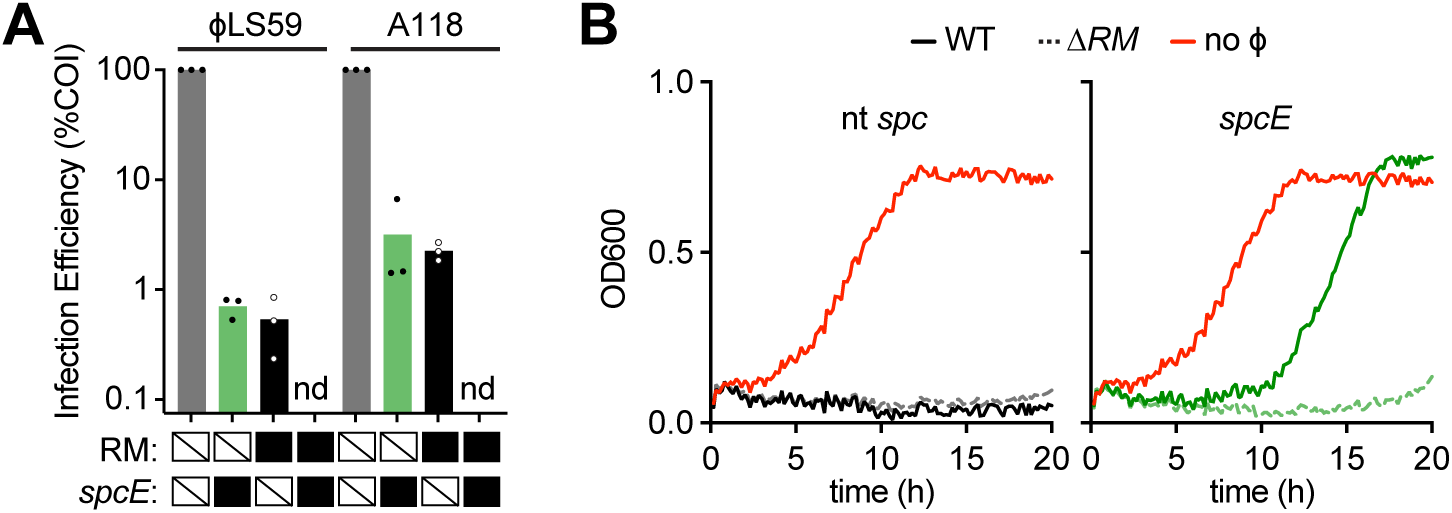
RM and type VI CRISPR systems synergize in anti-phage immunity. **(A)** Center of infection (COI) measurements for strains with (black boxes) or without (strikes) indicated defense. Cells were infected with the indicated phage at MOI 10 for 5 minutes, then washed 3 times before plating on a lawn of ΔRM cells lacking CRISPR. Infection efficiency reported displayed as a percentage of the number of PFU detected in the ΔRM non-targeting spacer infection. **(B)** Growth curves during A118 infection of LS1. Strains of the indicated genotype were infected at MOI=10, OD600=0.01. OD600 was monitored for 20 hours after infection. Solid lines indicate wild-type, RM^+^ strains, dashed lines indicate ΔRM strains. Uninfected controls are shown in red. The mean of 3 biological replicates is plotted.

Next, we wondered whether Cas13 might provide a more pronounced benefit during infection by phages that are less RM-sensitive. Of the phages we examined, ϕLS59 is by far the most RM-sensitive, with 21 RM recognition sites in its genome. Therefore, we examined infection efficiency and cell growth during infection by the slightly more RM- resistant phage A118, which only has 9 RM sites in its genome. We confirmed that A118 is sensitive to Cas13 targeting with a spacer targeting an early lytic gene (*spcE*) (Fig S5). As expected, we observed a higher rate of RM escape for A118 in the center of infection assay (Fig 5A). As with ϕLS59, A118 was unable to productively infect cells equipped with both RM and Cas13 immunity. Next, we monitored growth in liquid culture by optical density during A118 infection, as in Fig 3C. In contrast to the unperturbed growth we observed for RM+ cells during ϕLS59 infection (Fig 3C), the same strain rapidly succumbed to A118 infection (Fig 5B). The strain lacking RM but armed with *spcE* exhibited a prolonged growth defect during A118 infection. However, the strain with both RM and *spcE* began to recover from infection after 10 hours. The enhanced survival of this strain indicates that Cas13 confers a protective advantage to bacteria infected by phage whose sensitivity to RM interference is incomplete.

## DISCUSSION

Here we have investigated the fates of bacteria that simultaneously use abortive infection and cell-autonomous immunity to combat phage infection. Previous reports have established that type VI CRISPR systems unleash non-specific RNA degradation activity upon recognition of targeted phage transcripts, which leads to growth cessation^13, 14^. Thus, type VI CRISPR immunity was thought to exclusively act through an abortive infection mechanism in which the infected cell does not recover, but restricts the propagation of phage, protecting uninfected cells in the greater population. In this study, we have shown that the natural type VI CRISPR host *Listeria seeligeri* interferes with phage infection using both Cas13 and RM systems. Under these circumstances, Cas13 activity and cell dormancy can be triggered while phage genome cleavage by RM is still underway. When this happens, the cell is not irreversibly committed to dormancy. Instead, the phage genome is cleared from the cell, removing the source of target transcripts, and the cell resumes growth. Our data further indicate that cells armed with both immune systems exhibit stronger defense against phage infection. We conclude that, if paired with a DNA-targeting effector, type VI CRISPR systems can provide non-abortive immunity.

Structural analyses of Cas13 homologs have revealed that duplex formation between the crRNA and target RNA induce conformational changes in the Cas13 protein, which bring its two HEPN nuclease domains into close proximity for catalysis^29^. While RM- mediated cleavage of phage genomes removes the source of target transcripts, there is no evidence to suggest that RM removes target transcripts, or deactivates Cas13 that has already engaged with target RNA. But our observation that cells recover from dormancy once phage DNA is eliminated implies that Cas13 might have the ability to deactivate its RNA degradation activity. One possible mechanism is that on-target *cis-* cleavage of target RNA allows Cas13 to reverse the structural changes associated with its active state. A similar model has been proposed as a deactivation mechanism for type III CRISPR systems, which cleave target RNA in a sequence-specific manner via the Csm3 RNase subunit of the Csm complex^30–33^. This cleavage is thought to shut down the downstream non-specific RNase and DNase activities associated with active type III CRISPR immunity, which would otherwise be detrimental to the host.

Alternatively, the *trans-*cleavage activity of two target-bound Cas13 monomers might mutually cleave the target RNA occupying the other monomer. It is also possible that target RNA simply disassociates from Cas13 at some rate, restoring the inactive state. Finally, negative regulatory factors might deactivate Cas13 by forcing target disengagement or by sequestering it in a subcellular location where damage is mitigated.

The combination of DNA cleavage and non-specific RNA degradation as immune effector activities also has parallels in type III CRISPR systems, which also sense RNA targets^30–33^. When the type III-A CRISPR-associated Csm complex engages with target RNA, four enzymatic activities are triggered. The Csm3 subunit catalyzes sequence-specific cleavage of the target RNA, the HD domain of Cas10 cleaves the transcribed target DNA, and the Palm domain of Cas10 synthesizes cyclic oligoadenylate signaling molecules which activate the non-specific RNase Csm6^30–36^. Similar to the findings described here, the DNase activity of Cas10 enables cell survival during infection and Csm6 RNA cleavage; cells with a mutation in the DNase domain of Cas10 exhibit Csm6-dependent growth arrest^37^. While the DNase activity of Cas10 is sufficient for anti-phage immunity when early-expressed targets are sensed, the nonspecific RNase activity of Csm6 is critical for immunity during targeting of late-expressed genes, or when mismatches between crRNA and target limit the extent of Csm activation^38^. In these situations, nonspecific RNA cleavage is thought to provide time for Cas10 to cleave DNA. Our observation that Cas13 activation enhances the elimination of phage DNA by RM systems (Fig 4E-F) suggests that it might play a similar role during type VI CRISPR immunity.

With few exceptions, most CRISPR systems directly destroy targeted viral DNA, enabling survival of the infected cell^4, 39^. Thus, when these systems acquire a new spacer from a previously unencountered invader, it is immediately subject to positive selection. In contrast, type VI CRISPR immunity induces cell dormancy, therefore newly acquired spacers would be subject to strong negative selection during infection.

Therefore, the finding that RM systems enable cells to escape dormancy provides a possible explanation for how new spacers are inherited by the offspring of cells that survived infection. Furthermore, it has been observed that RM cleavage of phage genomes strongly stimulates the acquisition of new spacers by generating free DNA ends, which act as substrates for the Cas1-2 spacer capture machinery^22, 40^. While the mechanism of spacer acquisition is not known for type VI systems, it could similarly be stimulated by RM interference.

Finally, a multitude of different anti-phage defenses beyond type VI CRISPR have been reported to operate via abortive infection^41^. These systems use diverse mechanisms to sense phage infection and respond by inducing growth arrest or cell death. For example, the *toxIN* toxin-antitoxin system of gram-negative bacteria induces endoribonucleolytic cleavage of host and phage transcripts in response to shutdown of host transcription during phage infection^42, 43^. While some abortive infection systems do result in cell death, our results raise the possibility that cells using nonlethal abortive infection mechanisms might also resume growth when working in concert with additional defenses. Whether cells have the capacity to reverse the activities of different abortive infection systems likely depends on the infection-associated signals responsible for triggering the collateral activities of immune effectors, and the ability of those signals to be resolved by other anti-phage defenses.

## ACKNOWLEDGEMENTS

We are grateful to all members of the Meeske lab for advice and encouragement, members of the Guo and Mitchell labs for helpful discussions, Richard Calendar (UC Berkeley) for providing A118 and U153 phages, and David Rudner (Harvard Medical School) / Xindan Wang (Indiana University) for providing plasmid templates pWX510 and pWX570. Support for this work comes from the NIH [R35GM142460, S10OD026741] and the University of Washington Royalty Research Foundation.

## AUTHOR CONTRIBUTIONS

The project was conceived by MCW, AER, and AJM. Experiments were performed by MCW, AER, and AJM. Bioinformatic analysis was conducted by MCW, SRM, and AJM. JL and MW isolated *L. seeligeri* strains. Pilot microscopy experiments were performed with assistance from ERR. The manuscript was written and edited by MCW, AER, SRM, and AJM.

## DECLARATION OF INTERESTS

The authors have no conflicts of interest.

---

**Figure S1.**
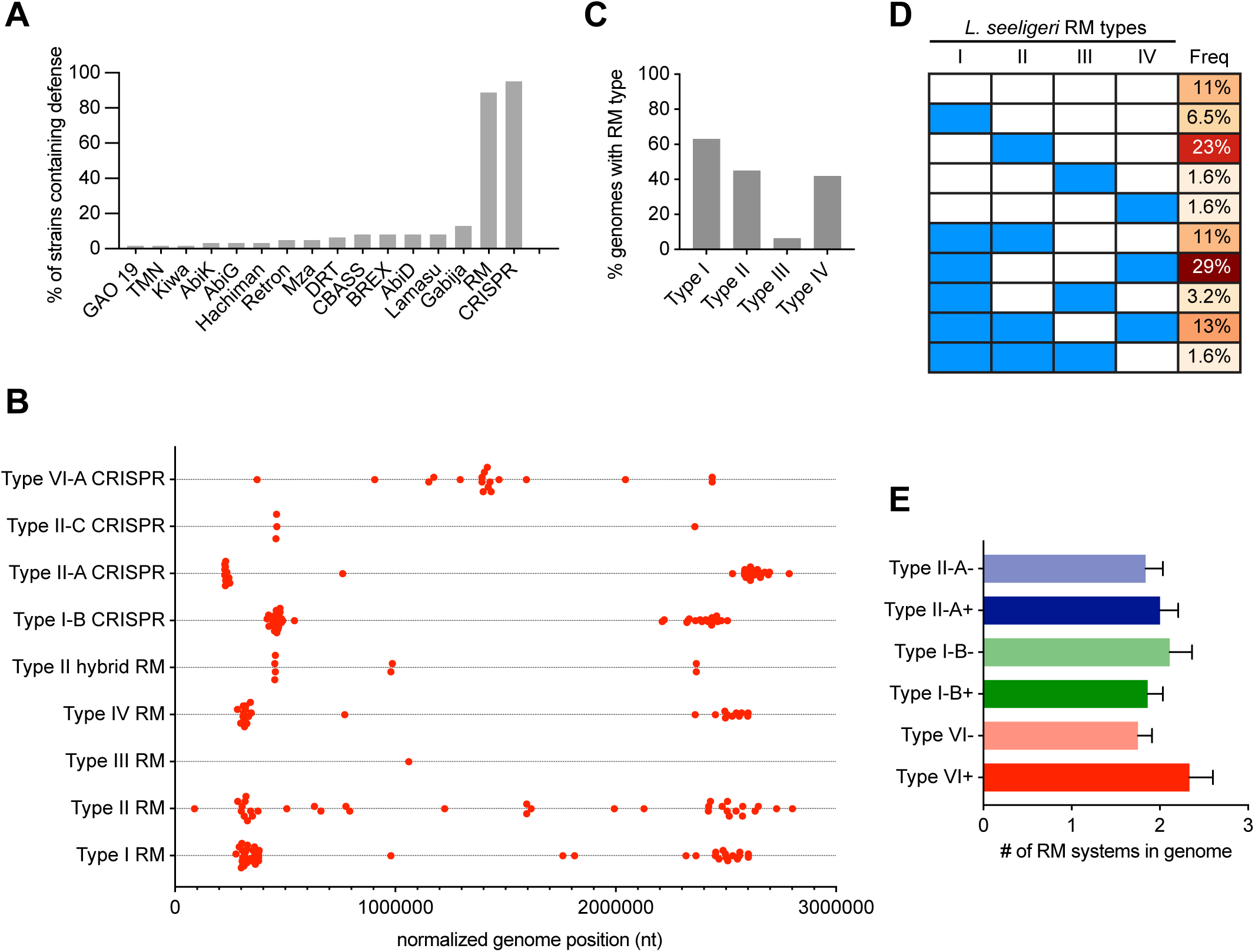
Bioinformatic analysis of anti-phage defenses in *L. seeligeri.* **(A)** Anti-phage defense systems detected in 62 *L. seeligeri* strains using PADLOC. **(B)** Genomic locations of CRISPR and RM systems within all 62 *L. seeligeri* genomes. Each genomic location was normalized to the position of the well-conserved, origin-proximal *dnaA* gene. **(C)** Percentage of 62 *L. seeligeri* genomes harboring the indicated RM type. **(D)** Combinations of RM types observed in 62 *L. seeligeri* genomes. Filled blue rectangles indicate the presence of the indicated RM type. **(E)** Tally of RM systems per genome in *Listeria* strains with or without the indicated CRISPR type. Error bars denote standard error of the mean.

**Figure S2.**
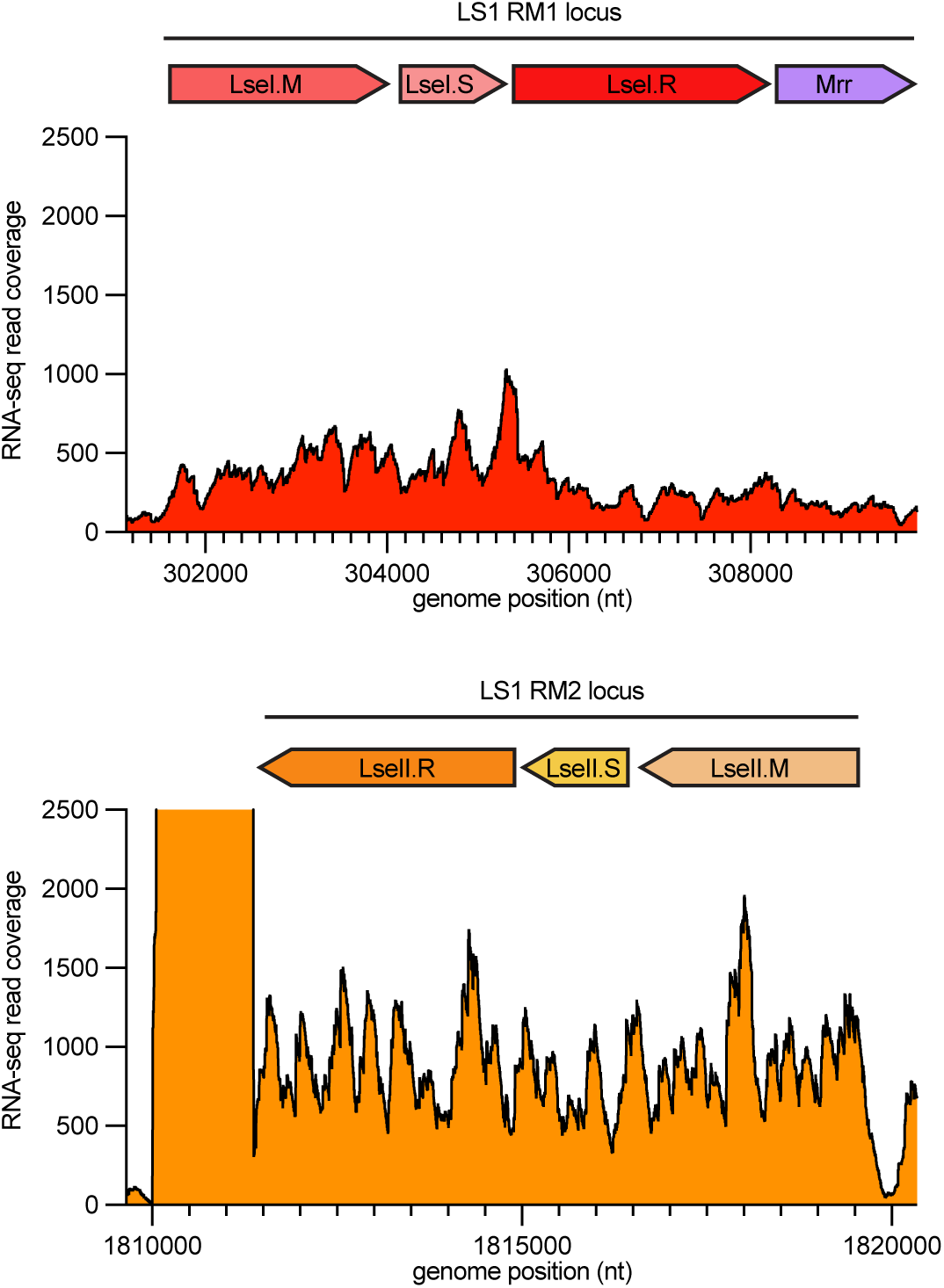
Both type I RM systems are expressed in *L. seeligeri* LS1. Diagrams of type I RM loci in the LS1 genome (methylase denoted M, specificity subunit denoted S, endonuclease denoted R, one locus also has a type IV nuclease denoted Mrr). RNA-seq read coverage at each nucleotide position of the two loci are plotted. The y-axis reflects the number of mapped reads overlapping each nucleotide position.

**Figure S3.**
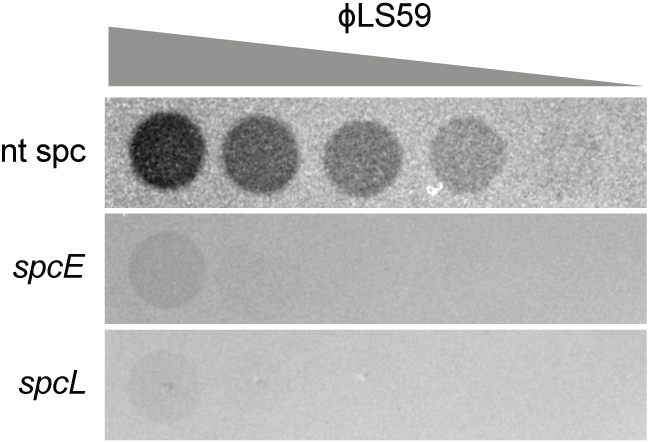
ϕLS59 is sensitive to type VI CRISPR immunity. Plaque assay of ϕLS59 on ΔRM strains harboring either a non-targeting spacer, spacer targeting early lytic genes (*spcE*) or spacer targeting late lytic genes (*spcL*). Serial tenfold dilutions were made from the phage stock, and 2µL of each dilution was spotted on the indicated bacterial lawn.

**Figure S4.**
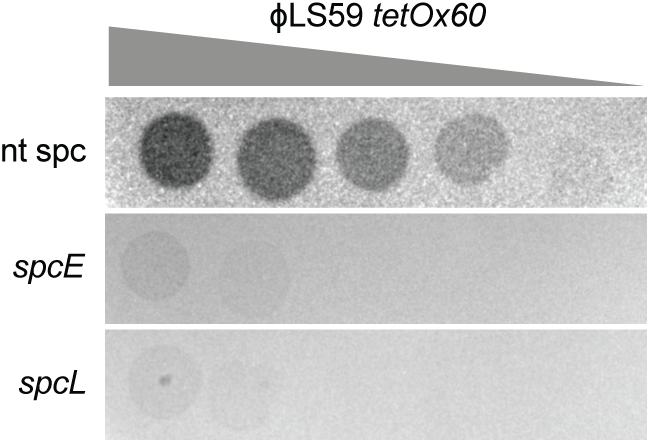
ϕLS59 tetOx60 is viable and sensitive to type VI CRISPR immunity. Plaque assay of ϕLS59 *tetOx60* on ΔRM strains harboring either a non-targeting spacer, spacer targeting early lytic genes (*spcE*) or spacer targeting late lytic genes (*spcL*). Serial tenfold dilutions were made from the phage stock, and 2µL of each dilution was spotted on the indicated bacterial lawn.

**Figure S5.**
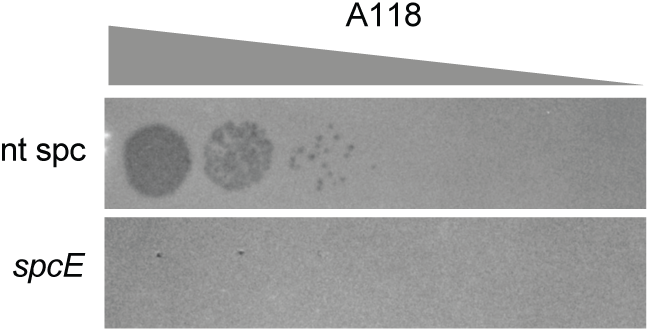
Phage A118 is sensitive to type VI CRISPR immunity. Plaque assay of A118 on ΔRM strains harboring either a non-targeting spacer or spacer targeting early lytic genes (*spcE*). Serial tenfold dilutions were made from the phage stock, and 2µL of each dilution was spotted on the indicated bacterial lawn.

## METHODS

### Data availability

All *L. seeligeri* genome sequences reported here have been annotated and uploaded to NCBI through Genbank. Lists of strains, plasmids, and oligonucleotides used in this study are available in Table S1. Further information and requests for resources and reagents should be directed to and will be fulfilled by the corresponding author, Alexander Meeske.

### Bacterial strains and growth conditions

*L. seeligeri* strains were propagated in Brain Heart Infusion (BHI) broth or agar at 30°C. Where appropriate, BHI was supplemented with the following antibiotics for selection: nalidixic acid (50 µg/mL) chloramphenicol (10 µg/mL), erythromycin (1 µg/mL), or kanamycin (50 µg/mL). For cloning, plasmid preparation, and conjugative plasmid transfer, *E. coli* strains were cultured in Lysogeny Broth (LB) medium at 37°C. Where appropriate, LB was supplemented with the following antibiotics: ampicillin (100 µg/mL), chloramphenicol (25 µg/mL), kanamycin (50 µg/mL). For conjugative transfer of *E. coli – Listeria* shuttle vectors, plasmids were purified from Turbo Competent *E. coli* (New England Biolabs) and transformed into the *E. coli* conjugative donor strains SM10 λ*pir* or S17 λ*pir* ^44^.

### Phage propagation

Unless otherwise stated, all phage infections were performed in BHI medium supplemented with 5mM CaCl2. To generate phage lysates, existing phage stocks were diluted to single plaques on a lawn of *L. seeligeri* LS1 *ΔRM1 ΔRM2*, and a single plaque was purified twice to ensure homogeneity. 5 mL of cell culture was infected with phage at MOI 0.1, OD 0.1 and the infection proceeded overnight. The lysate was centrifuged for 20 minutes at 4,000 rpm, then the supernatant was filtered using a 0.45 µm pore syringe filter.

### Plasmid construction and preparation

All genetic constructs for expression in *L. seeligeri* were cloned into the following three compatible shuttle vectors, each of which contains an origin of transfer sequence for mobilization by transfer genes of the IncP-type plasmid RP4. These transfer genes are integrated into the genome of the *E. coli* conjugative donor strains SM10 λ*pir* and S-17 λ*pir* ^44^. All plasmids used in this study, along with details of their construction, can be found in Table S1.

pPL2e – single-copy plasmid conferring erythromycin resistance that integrates into the *tRNA^Arg^* locus in the *L. seeligeri* chromosome ^45^.

pAM8 – *E. coli – Listeria* shuttle vector conferring chloramphenicol resistance ^24^. pAM326 - *E. coli – Listeria* shuttle vector conferring kanamycin resistance^17^.

### Center of infection assay

1. *L. seeligeri* strains were grown to mid-exponential phase, and 0.5 mL of culture at OD600 0.5 was infected with phage or derivatives at the indicated MOI (10 for most experiments). Adsorption was carried out for 20 minutes. To remove unbound phage, the cell suspension was washed three times with 1 mL BHI, and was transferred to a fresh tube during the third wash. The samples were resuspended to a final volume of 1 mL. Infective centers were enumerated by titering PFU on a lawn of naïve *L. seeligeri* LS1 *ΔRM1 ΔRM2*.
2. *E. coli – L. seeligeri* conjugation:

All genetic constructs for expression in *L. seeligeri* were introduced by conjugation with the *E. coli* donor strains SM10 λ*pir*, S-17 λ*pir* ^44^, or for allelic exchange (see below), β2163 Δ*dapA* ^46^. Donor cultures were grown overnight in LB medium supplemented with the appropriate antibiotic (25 µg/mL chloramphenicol for pPL2e-derived plasmids, 100 µg/mL ampicillin for pAM8-derived plasmids, or 50 µg/mL kanamycin for pAM326- derived plasmids) at 37°C. Recipient cultures were grown overnight in BHI medium supplemented with the appropriate antibiotic (1 µg/mL erythromycin for pPL2e-derived plasmids, 10 µg/mL chloramphenicol for pAM8-derived plasmids, 50 µg/mL kanamycin for pAM326-derived plasmids) at 30°C. 100 µL each of donor and recipient culture were diluted into 10 mL of BHI medium, and concentrated onto a filter disc (Millipore-Sigma, HAWP04700) using vacuum filtration. Filter discs were laid onto BHI agar supplemented with 8 µg/mL oxacillin (which weakens the cell wall and enhances conjugation) and incubated at 37°C for 4 hr. Discs were removed, cells were resuspended in 2 mL BHI, and transconjugants were selected on medium containing 50 µg/mL nalidixic acid (which kills donor *E. coli* but not recipient *L. seeligeri*) in addition to the appropriate antibiotic for plasmid selection. Transconjugants were isolated after 2-3 days incubation at 30°C.

### Gene deletions and replacements in *Listeria*

Allelic exchange plasmids were generated by cloning 1kb homology arms flanking the genomic region to be deleted into the suicide vector pAM215 ^14^, which does not replicate in *Listeria*, and contains a chloramphenicol resistance cassette and *lacZ* from *Geobacillus stearothermophilus*. These plasmids were then transformed into the *E. coli* donor strain β2163 Δ*dapA*^46^, which is auxotrophic for diaminopimelic acid (DAP), selecting on LB medium supplemented with the appropriate antibiotic and 1.2 mM DAP. Conjugation was carried out as described above, except all steps were carried out in the presence of 1.2 mM DAP. Transconjugants were selected on media lacking DAP and containing 50 µg/mL nalidixic acid, to ensure complete killing of donor *E. coli*, as well as 10 µg/mL chloramphenicol to select for integration of the pAM215-derived plasmid.

Chloramphenicol-resistant colonies were patched on BHI supplemented with 100 µg/mL 5-Bromo-4-Chloro-3-Indolyl β-D-Galactopyranoside (X-gal) and confirmed *lacZ*+ by checking for blue colony color. Plasmid integrants were passaged 3-4 times in BHI at 30° in the absence of antibiotic selection, to permit loss of the integrated plasmid.

Cultures were screened for plasmid excision by dilution and plating on BHI + X-gal. White colonies were checked for chloramphenicol sensitivity, then chromosomal DNA was prepared from each, and tested for the desired deletion by PCR using primers flanking the deletion site. Deletions were confirmed by Sanger sequencing.

### Type VI spacer cloning and expression in *L. seeligeri*

All spacers used in this study were cloned into entry vector pAM305, which is a site-specific chromosomally integrating vector containing a type VI repeat-spacer-repeat miniarray, expressed from the native promoter, with BsaI restriction sites in the spacer allowing for facile replacement. All strains used in this study have the native 5 spacer array deleted from the type VI CRISPR locus (with the promoter and *cas13* gene left intact). Spacer constructs were integrated into the genome via conjugation and expressed as the sole type VI spacer.

### Bacterial genome sequencing and assembly

Chromosomal DNA was prepared from each *L. seeligeri* isolate by lysozyme digestion of the cell wall, followed by cell lysis with 1% sarkosyl, then phenol-chloroform extraction and ethanol precipitation. 1 ng of chromosomal DNA was used to make an NGS library using the Illumina Nextera XT DNA Library Preparation Kit according to the manufacturer’s instructions. Library quality was confirmed by analysis on Agilent TapeStation, then 2x150bp paired-end sequencing was carried out on the Illumina NextSeq platform. Raw reads were quality-trimmed using Sickle (https://github.com/najoshi/sickle) using a quality cutoff of 30 and length cutoff of 45.

Trimmed reads were assembled using SPAdes (http://cab.spbu.ru/software/spades/) with the default parameters, which resulted in assembled contigs. These contigs were mapped onto the completed reference genome of *L. seeligeri* SLCC3954 using Medusa (http://combo.dbe.unifi.it/medusa/) with the default parameters, generating scaffold assemblies. In each draft genome assembly, only one scaffold represented the fully assembled genome.

### Construction of ϕLS59 tetOx60

The tetOx60 array was inserted into the ϕLS59 genome by homologous recombination and Cas9 selection. A recombination template plasmid (pAM542) was generated containing the tetOx60 array flanked on each side by 500 bp of ϕLS59 sequence. The insertion site was downstream of the late lytic genes, in a putative accessory region of the genome. The insertion site lay adjacent to a PAM for SpyCas9, which we mutated in the recombination template to confer Cas9 resistance to recombinants. The recombination template plasmid pAM542 was introduced into LS1 ΔRM, this strain was infected with ϕLS59 in BHI top agar (allowing recombinants to be generated), and a phage stock was harvested. A Cas9 spacer targeting the site disrupted by tetOx60 insertion was cloned into pAM307 to generate pAM545 and introduced into LS1 ΔRM. The ϕLS59 stock passaged on LS1 ΔRM carrying the pAM542 repair template was used to infect LS1 ΔRM carrying pAM545, and Cas9-resistant escaper mutants were isolated. Two mutant phage isolates were Sanger sequenced across the tetOx60 insertion site, and found to contain the precise insertion.

### Microscopy

Timelapse imaging of phage infection was performed with mid-exponential phase cells, infected at OD600 of 0.01, MOI of 10, in BHI medium supplemented with 2mM CaCl2. Adsorption was allowed to occur for 10 minutes, then cells were loaded into microfluidic chambers using the CellASIC ONIX2 Microfluidic System (Millipore-Sigma). After cells became trapped in the chamber, they were supplied with BHI medium under a constant flow of 5 µL per hr. Phase contrast images were captured at 1000x magnification every 10 minutes for 5 hours after infection, using a Nikon Ti2e inverted microscope equipped with a Hamamatsu Orca-Fusion SCMOS camera and temperature-controlled enclosure set to 30°C. SYTOX Green stain was imaged using a GFP filter set, and TetR-mCherry signal with Texas Red filter set, both using an Excelitas Xylis LED Illuminator set to 6% power with an exposure time of 300 ms. Timelapse images were aligned and processed using NIS Elements software. Quantitative analysis of cell fates and phage genome foci were performed in Fiji.

### Bioinformatics

CRISPR-cas loci and RM loci were identified in *Listeria* genomes via TBLASTN searches of the 62 newly sequenced *L. seeligeri* genomes, as well as all *Listeriaceae* in the NCBI ‘wgs’ database (excluding *L. monocytogenes*). Cas proteins from each CRISPR type were used as queries for CRISPR loci, with a E-value cutoff of 10^-4^. For RM loci, all unique *Listeria* RM systems available on REBASE (rebase.neb.com) were used as queries in the TBLASTN searches above. Non-CRISPR anti-phage defense systems of the 62 *L. seeligeri* genomes were identified using PADLOC^47^; only systems containing all required genes are reported. To normalize genome positions, *dnaA* homologs were identified in each genome, and defense positions in Fig S1 are reported relative to each *dnaA* gene.

### CFU survival assay

Mid-exponential phase cells were diluted to OD600 of 0.005 in BHI medium supplemented with 5mM CaCl2, and a pre-infection sample was harvested for CFU enumeration. Cells were infected with phage at MOI 10, and additional CFU samples were harvested at 0.5, 4, and 24h post-infection. 8 ten-fold serial dilutions in BHI were made from culture samples at the time of harvest, and 5 µL of each dilution was spotted on BHI agar plates supplemented with 50mM sodium citrate, to prevent additional infections from occurring on the plate. Viable colonies were counted after 2 days incubation at 30°C.

### Methylation site identification

Genome-wide methylation motifs on LS1 DNA were identified by PacBio sequencing of genomic DNA harvested from LS1 ΔRM1 (to identify the RM2 methylation motif) and LS1 ΔRM2 (to identify the RM1 methylation motif). Sequencing was performed by CD Genomics. Methylation sites were identified using the BaseMod Analysis tool in the SMRT Analysis package, with assistance from the University of Washington PacBio Sequencing Center.

### RNA isolation and Northern blot analysis

12ml cultures of *L. seeligeri* strains were infected with ϕLS59 at MOI 10 and OD600 of 0.1. At indicated time points post-infection, 1 mL aliquots of culture were pelleted by centrifugation at 8,000 rpm at 4°C, and frozen. Cell pellets were resuspended in 80 µL RNase-free PBS, and lysed by a 5 min treatment with lysozyme at 20 µg/mL, followed by the addition of 1% sarkosyl. RNA was purified from these lysates by adding 300 µL Trizol LS (Thermo), followed by 80 µL of chloroform. Samples were centrifuged at 15,000 rpm for 15 minutes, and RNA was precipitated from the upper aqueous phase by the addition of glycoBlue co-precipitate and 200 µL of isopropanol. RNA pellets were washed with 500 µL of 80% ethanol, air-dried, and resuspended in RNase-free water. For Northern blot analysis, 1 µg RNA per sample was diluted in RNA loading dye (95% formamide, 1.8 mM EDTA, 0.5% bromophenol blue), heated at 95°C for 3 minutes, cooled on ice for 1 minute, and separated by gel electrophoresis on a precast 15% polyacrylamide denaturing TBE-urea gel. RNA was transferred to an Invitrogen Immobilon Ny+ nylon membrane using a Bio-Rad Mini Trans-Blot Cell filled with 1X TBE. RNA was fixed to membranes using a Analytik Jena UV crosslinker (Optimal Crosslinking setting). Membranes were pre-hybridized 30 min at 42°C in 2X SSC containing 1% SDS. 250 pmol Cy3-labeled ssDNA probe complementary to the 5S rRNA (oAM734) was hybridized to the membrane overnight at 42°C. Membranes were washed twice with 2X SSC 0.1% SDS, once with 1X SSC 0.1% SDS, then imaged using an Azure Sapphire Biomolecular Imager.

## Notes

### Competing Interest Statement

The authors have declared no competing interest.

