## Supplementary File 1 for "Phage genome cleavage enables resuscitation from Cas13-induced bacterial dormancy"

SMRT Analysis / Analysis Results

### BaseMod - DeltaRM1

**SUCCESSFUL** Copy Delete

- **Analysis Overview**
  - Status
  - Thumbnails
  - Display All
- Mapping Report
- Coverage
- Base Modifications
- Modified Base Motifs
- Data

#### Display All

##### Status

| | |
|---|---|
| **Analysis** | BaseMod - DeltaRM1 |
| **Analysis ID** | 7443 |
| **From Multi-Job** | 7442 |
| **Status** | SUCCESSFUL: 55 tasks finished |
| **Created By** | kmiyamot |
| **Date Created** | 2021-05-26, 10:35:27 AM |
| **Date Updated** | 2021-05-26, 11:10:18 AM |
| **Application** | Base Modification Analysis |
| **SMRT Link Version** | 10.1.0.119588 |
| **Inputs** | |
| **Path** | /net/eichler/vol28/projects/sequencing/pacbio/smrt-link/userdata/jobs_root/0000/0000007/0000007443 |

Analysis Parameters

##### Thumbnails

Mapped Polymerase Read Length

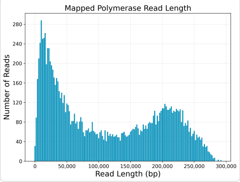


Coverage Across NC_013891.1.

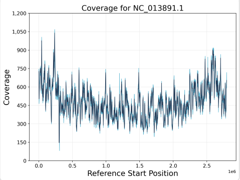


Kinetic Detections Histogram

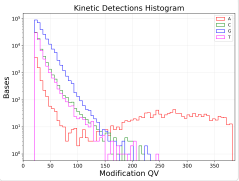


Alignment Length

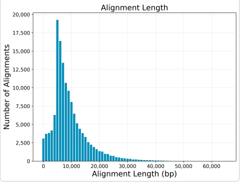


Depth of Coverage Distribution

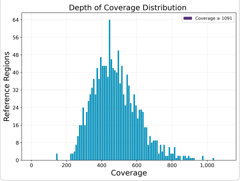


Modification QVs

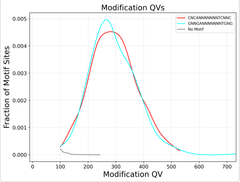


Alignment Concordance

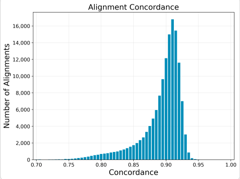


Coverage vs. [GC] Content

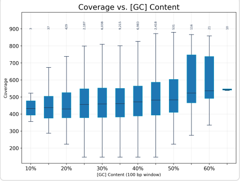


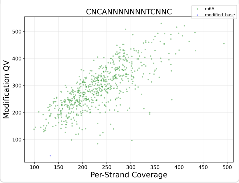


Mapped Concordance vs. Alignment Length

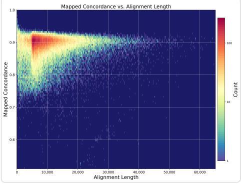


Per-Base Kinetic Detections

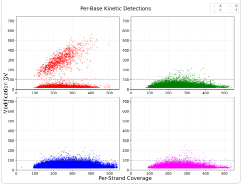


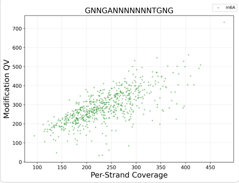


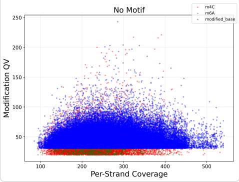

#### Mapping Report

| Value | Analysis Metric |
|---:|---|
| 96.67% | Percentage of Bases (mapped) |
| 134,696 | Number of Subreads (total) |
| 133,188 | Number of Subreads (mapped) |
| 1,508 | Number of Subreads (unmapped) |
| 98.88% | Percentage of Subreads (mapped) |
| 1.11% | Percentage of Subreads (unmapped) |
| 88.97% | Mean Concordance (mapped) |
| 1,367,910,858 | Number of Subread Bases (mapped) |
| 140,318 | Number of Alignments |
| 9,748 | Alignment Length Mean (mapped) |
| 11,580 | Alignment Length N50 (mapped) |
| 22,615 | Alignment Length 95% (mapped) |
| 65,015 | Alignment Length Max (mapped) |
| 12,433 | Number of Polymerase Reads (mapped) |
| 112,814 | Polymerase Read Length Mean (mapped) |
| 196,099 | Polymerase Read N50 (mapped) |
| 247,740 | Polymerase Read Length 95% (mapped) |
| 293,849 | Polymerase Read Length Max (mapped) |

#### Mapping Statistics Summary

| Sample | Movie | Number of Polymerase Reads (mapped) | Polymerase Read Length Mean (mapped) | Polymerase Read N50 (mapped) | Number of Subreads (mapped) | Number of Subread Bases (mapped) | Subread Length Mean (mapped) | Mea Conc (map |
|---|---|---|---|---|---|---|---|---|
| All Samples | All Movies | 12,433 | 112,814 | 196,099 | 133,188 | 1,367,910,858 | 9,748 | 88.97 |
| MC0513 MIX 383-5 | m64083_210514_071054 | 12,433 | 112,814 | 196,099 | 133,188 | 1,367,910,858 | 9,748 | 88.97 |

#### Mapped Polymerase Read Length

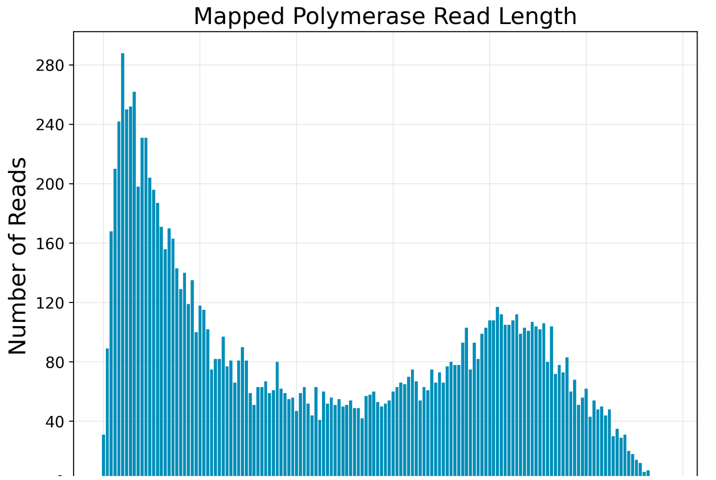

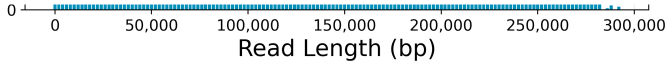


#### Alignment Length

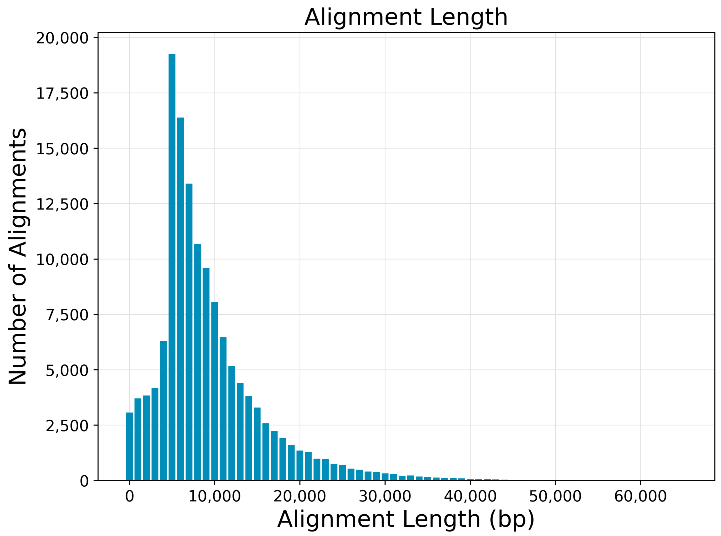


#### Alignment Concordance

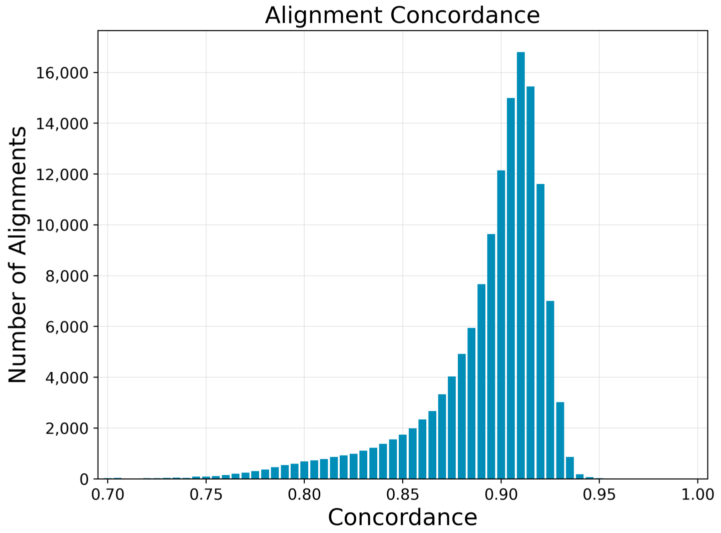


#### Mapped Concordance vs. Alignment Length

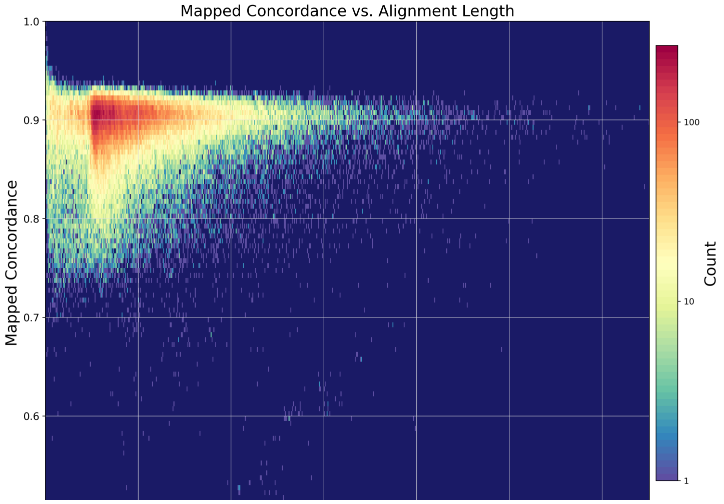

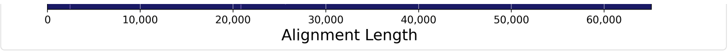


#### Coverage

| Value | Analysis Metric |
| --- | --- |
| 479 | Mean Coverage |
| 0.00% | Missing Bases |

#### Coverage Across Reference

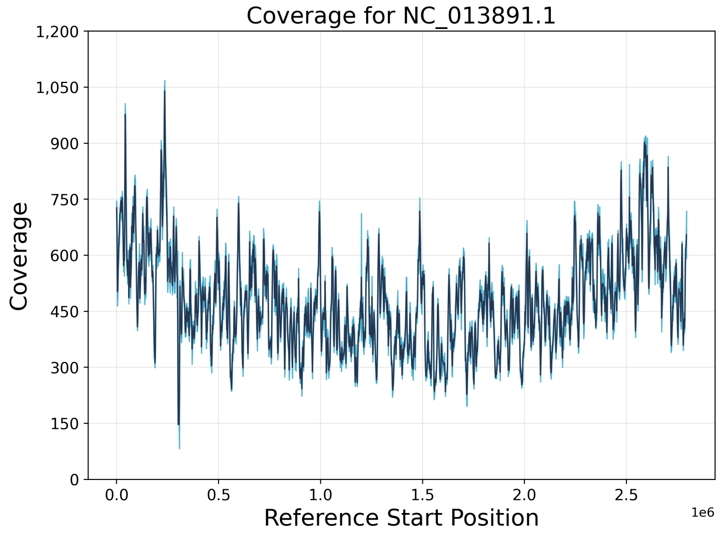


#### Depth of Coverage

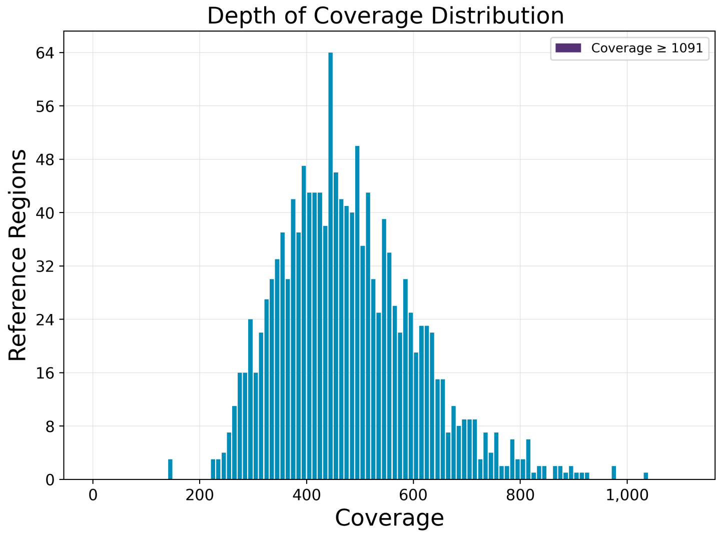


#### Coverage vs. [GC] Content

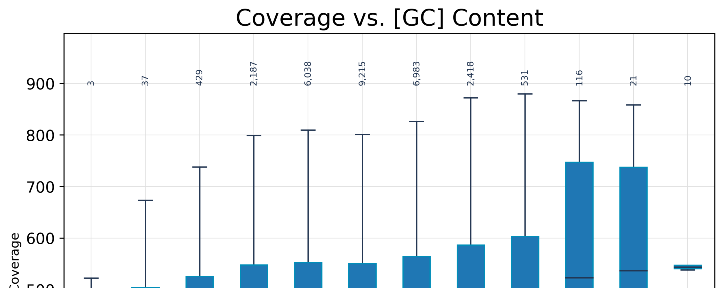

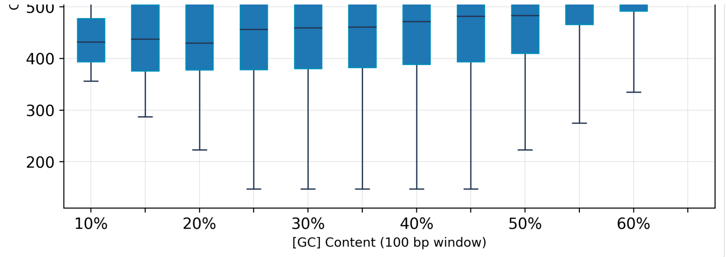


#### Kinetic Detections

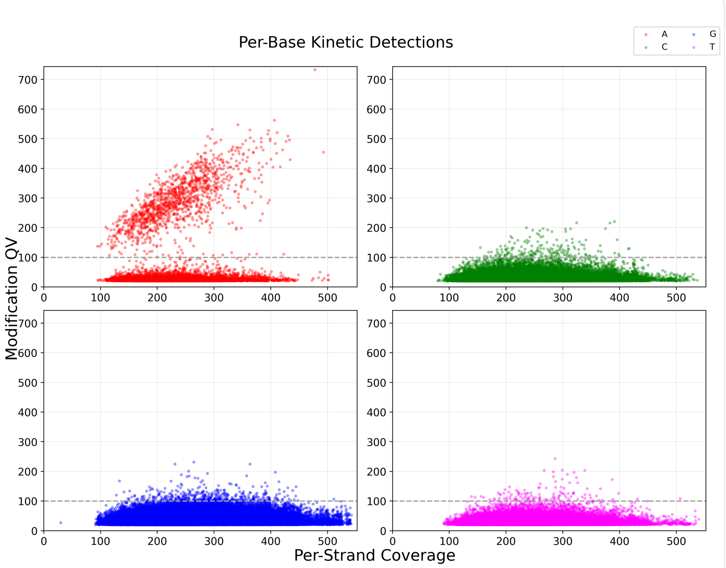


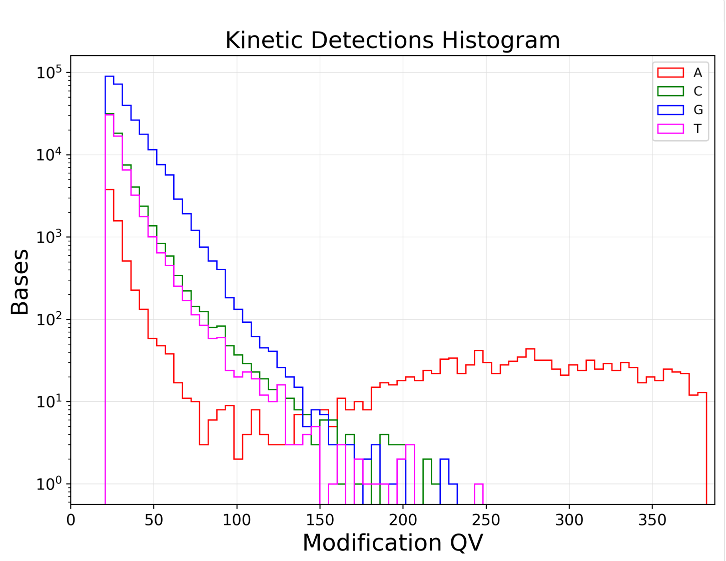


#### Modified Base Motifs

| Motif | Modified Position | Modification Type | % of Motifs Detected | # of Motifs Detected | # of Motifs in Genome | Mean QV | Mean Coverage | Partner Motif |
|---|---|---|---|---|---|---|---|---|
| CNCANNNNNNNTCNNC | 4 | m6A | 98.7% | 620 | 628 | 294.1 | 234.6 | GNNGANNNNN |
| GNNGANNNNNNNTGNG | 5 | m6A | 98.2% | 617 | 628 | 293.6 | 236.8 | CNCANNNNNN |

#### Modification QVs

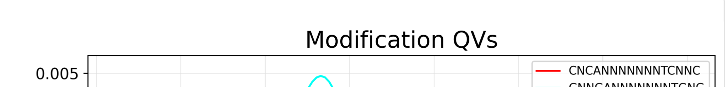

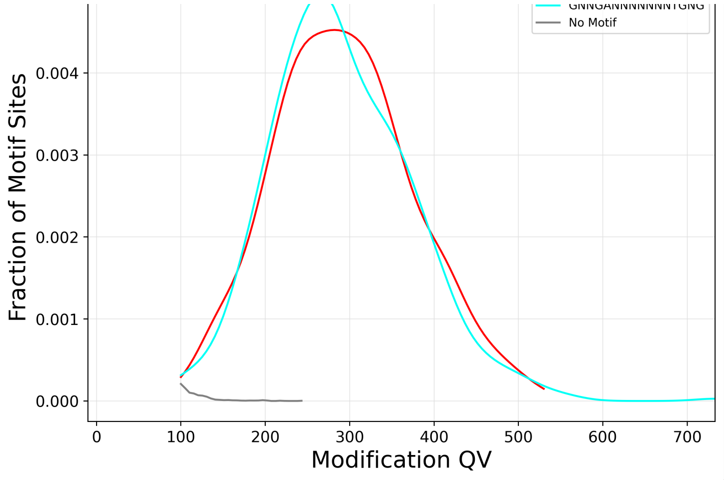


#### ModQV Versus Coverage By Motif

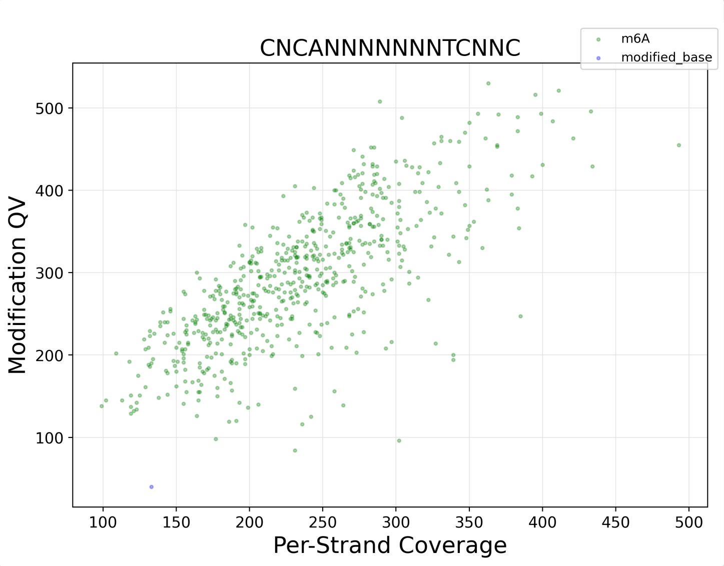


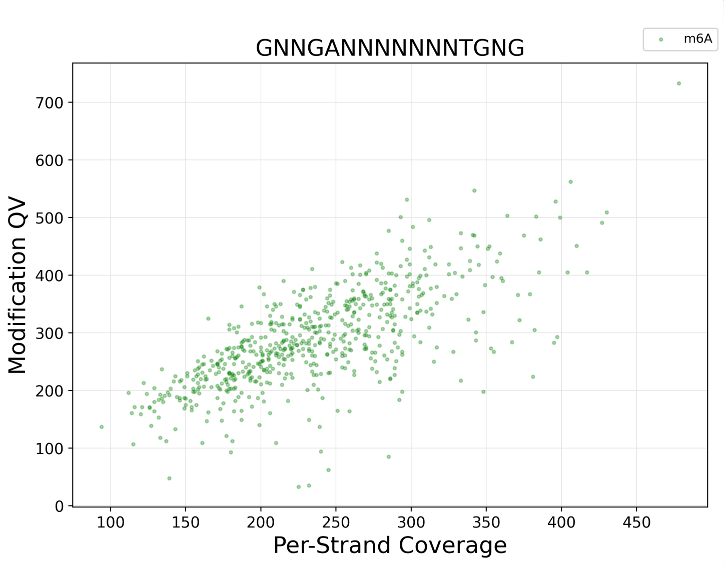


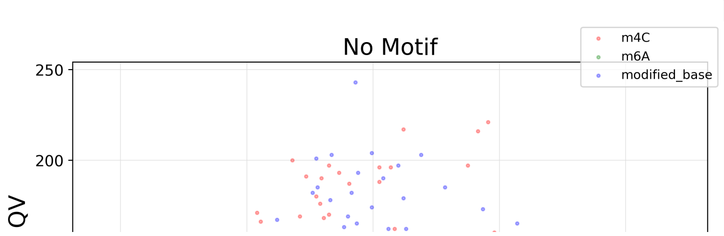

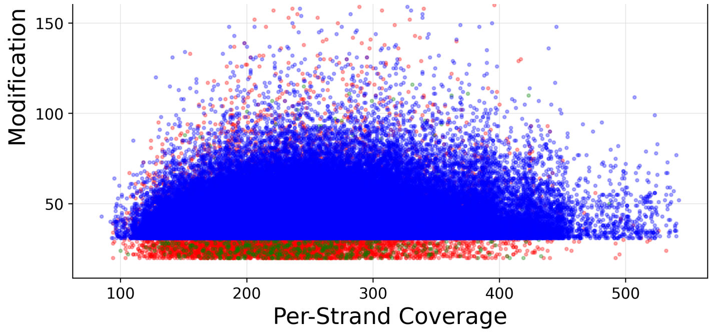


#### File Downloads

Edit Output File Name Prefix

**Example:**analysis-DeltaRM1-7443

#### IGV Visualization Files

| File | Path | Size | Type |
|---|---|---|---|
| Mapped BAM Index | http://eee-smrt.gs.washington.edu:9090/job-data/pb_basemods/7a651345-c054-41f8-ae02-18c55bac5311/call-auto_consolidate_alignments/execution/mapped.bam.bai | 250 KB | bam_bai |
| Mapped BAM | http://eee-smrt.gs.washington.edu:9090/job-data/pb_basemods/7a651345-c054-41f8-ae02-18c55bac5311/call-auto_consolidate_alignments/execution/mapped.bam | 3 GB | bam |
| Per-Base IPDs for IGV | http://eee-smrt.gs.washington.edu:9090/job-data/pb_basemods/7a651345-c054-41f8-ae02-18c55bac5311/call-gather_bigwig/execution/ipds.bw | 19 MB | bigwig |

##### SMRT Link Log

```
[INFO] [2021-05-26T10:35:32.473-07:00] Initial Setup of resources. Starting to run engine job id:7443 type:analysis Parent
MultiJobId:7442
[INFO] [2021-05-26T10:35:32.473-07:00] Job output directory is /net/eichler/vol28/projects/sequencing/pacbio/smrt-
link/userdata/jobs_root/0000/0000007/0000007443
[INFO] [2021-05-26T10:35:32.832-07:00] Final task options before converting to Cromwell inputs:
[INFO] [2021-05-26T10:35:32.834-07:00] [{
  "id": "run_find_motifs",
  "value": true,
  "optionTypeId": "boolean"
}, {
  "id": "consolidate_aligned_bam",
  "value": true,
  "optionTypeId": "boolean"
}, {
  "id": "dataset_filters",
  "value": "",
  "optionTypeId": "string"
}, {
  "id": "kineticstools_p_value",
  "value": 0.001,
  "optionTypeId": "float"
}, {
  "id": "kineticstools_compute_methyl_fraction",
  "value": true,
  "optionTypeId": "boolean"
}, {
  "id": "motif_min_score",
  "value": 100,
  "optionTypeId": "integer"
}, {
  "id": "motif_min_fraction",
  "value": 0.3,
  "optionTypeId": "float"
}, {
  "id": "mapping_min_concordance",
  "value": 70.0,
  "optionTypeId": "float"
}, {
  "id": "mapping_min_length",
  "value": 50,
  "optionTypeId": "integer"
}, {
  "id": "mapping_biosample_name",
  "value": "",
  "optionTypeId": "string"
}, {
  "id": "mapping_pbmm2_overrides",
  "value": "",
  "optionTypeId": "string"
}, {
  "id": "downsample_factor",
  "value": 0,
  "optionTypeId": "integer"
}, {
  "id": "kineticstools_identify_mods",
  "value": "m4C,m6A",
  "optionTypeId": "string"
}]
[INFO] [2021-05-26T10:35:32.835-07:00] Workflow options from job submission:
List(ServiceTaskIntOption(cromwell.workflow_options.max_nchunks,48,integer),
ServiceTaskStrOption(cromwell.engine_options.memory,32 G,string),
ServiceTaskIntOption(cromwell.workflow_options.nproc,8,integer),
ServiceTaskStrOption(cromwell.workflow_options.log_level,INFO,string),
ServiceTaskIntOption(cromwell.workflow_options.memory_per_core,4,integer),
ServiceTaskBooleanOption(cromwell.engine_options.read_from_cache,false,boolean),
ServiceTaskStrOption(cromwell.engine_options.backend,computecfg_00,string),
ServiceTaskBooleanOption(cromwell.engine_options.write_to_cache,true,boolean))
[INFO] [2021-05-26T10:35:32.836-07:00] Final engine options for analysis:
{"read_from_cache":false,"write_to_cache":true,"backend":"computecfg_00","default_runtime_attributes":
{"backend":"computecfg_00","maxRetries":1,"memory":"32 G","username":"kmiyamot"}}
[INFO] [2021-05-26T10:35:32.846-07:00] Submitting Cromwell workflow to http://localhost:9096
[INFO] [2021-05-26T10:35:32.894-07:00] Started Cromwell workflow 7a651345-c054-41f8-ae02-18c55bac5311 from SMRT Link job
c1d55765-7507-4f22-a3b2-f1193b426a40
[INFO] [2021-05-26T10:35:42.916-07:00] Cromwell workflow now in state SUBMITTED
[INFO] [2021-05-26T10:35:42.970-07:00] 0 tasks finished
[INFO] [2021-05-26T10:36:05.076-07:00] Cromwell workflow now in state RUNNING
[INFO] [2021-05-26T10:36:05.112-07:00] Created symlink to
/net/eichler/vol28/projects/sequencing/pacbio/nobackups/smrtlink_job_data/cromwell-executions/pb_basemods/7a651345-c054-41f8-
ae02-18c55bac5311
[INFO] [2021-05-26T10:36:05.131-07:00] 1 task finished, task pb_basemods.update_subreads and 0 others running
[INFO] [2021-05-26T10:36:37.000-07:00] 2 tasks finished, task pb_basemods.dataset_filter and 0 others running
[INFO] [2021-05-26T10:36:57.877-07:00] 3 tasks finished, task get_input_sizes.get_ref_size and 1 other running
[INFO] [2021-05-26T10:37:22.898-07:00] 5 tasks finished, task mapping.split_reads and 0 others running
[INFO] [2021-05-26T10:37:52.909-07:00] 6 tasks finished, task mapping.pbmm2_align and 5 others running
[INFO] [2021-05-26T10:40:03.935-07:00] 10 tasks finished, task mapping.pbmm2_align and 1 other running
[INFO] [2021-05-26T10:41:04.250-07:00] 12 tasks finished
[INFO] [2021-05-26T10:42:04.510-07:00] 13 tasks finished, task mapping.mapping_stats and 0 others running
[INFO] [2021-05-26T10:43:04.809-07:00] 16 tasks finished, task coverage_reports.summarize_coverage and 29 others running
[INFO] [2021-05-26T10:44:05.197-07:00] 18 tasks finished, task pb_basemods.auto_consolidate_alignments and 28 others running
[INFO] [2021-05-26T10:47:05.984-07:00] 19 tasks finished, task pb_basemods.ipdsummary and 27 others running
[INFO] [2021-05-26T10:49:06.765-07:00] 21 tasks finished, task pb_basemods.ipdsummary and 25 others running
[INFO] [2021-05-26T10:50:07.385-07:00] 26 tasks finished, task pb_basemods.ipdsummary and 20 others running
[INFO] [2021-05-26T10:51:07.793-07:00] 36 tasks finished, task pb_basemods.ipdsummary and 10 others running
```

```
[INFO] [2021-05-26T10:52:08.666-07:00] 43 tasks finished, task pb_basemods.ipdsummary and 3 others running
[INFO] [2021-05-26T10:53:09.289-07:00] 46 tasks finished, task pb_basemods.ipdsummary and 0 others running
[INFO] [2021-05-26T10:54:09.759-07:00] 47 tasks finished, task pb_basemods.gather_bigwig and 2 others running
[INFO] [2021-05-26T10:55:10.501-07:00] 51 tasks finished, task pb_basemods.find_motifs and 1 other running
[INFO] [2021-05-26T10:56:11.169-07:00] 52 tasks finished, task pb_basemods.find_motifs and 0 others running
[INFO] [2021-05-26T11:08:16.112-07:00] 54 tasks finished, task pb_basemods.motifs_report and 0 others running
[INFO] [2021-05-26T11:10:16.915-07:00] Cromwell workflow now in state SUCCEEDED
[INFO] [2021-05-26T11:10:17.219-07:00] 55 tasks finished
[INFO] [2021-05-26T11:10:18.580-07:00] Cromwell workflow completed, importing 13 outputs
[INFO] [2021-05-26T11:10:18.653-07:00] about to write datastore for job c1d55765-7507-4f22-a3b2-f1193b426a40
[INFO] [2021-05-26T11:10:18.696-07:00] Done writing datastore for job c1d55765-7507-4f22-a3b2-f1193b426a40
[INFO] [2021-05-26T11:10:18.696-07:00] Successfully completed running core job id:c1d55765-7507-4f22-a3b2-f1193b426a40
[INFO] [2021-05-26T11:10:18.902-07:00] Imported 11 files
Linked 4 reports to 3 datasets
Job id:7443 DataStoreFile 13381f58-08ec-2dd4-b6c5-21bce641491c imported
[INFO] [2021-05-26T11:10:18.918-07:00] Updated job c1d55765-7507-4f22-a3b2-f1193b426a40 state to SUCCESSFUL
[INFO] [2021-05-26T11:10:18.927-07:00] Job 7443 type:analysis state:SUCCESSFUL is NOT eligible for email sending. no
createdByEmail user. Skipping sending Email
```
