## Supplementary File 2 for "Phage genome cleavage enables resuscitation from Cas13-induced bacterial dormancy"

SMRT Analysis / Analysis Results

### BaseMod - DeltaRM2

**SUCCESSFUL** Copy Delete

- **Analysis Overview**
  - Status
  - Thumbnails
  - Display All
- Mapping Report
- Coverage
- Base Modifications
- Modified Base Motifs
- Data

#### Display All

##### Status

| | |
|---|---|
| **Analysis** | BaseMod - DeltaRM2 |
| **Analysis ID** | 7444 |
| **From Multi-Job** | 7442 |
| **Status** | SUCCESSFUL: 48 tasks finished |
| **Created By** | kmiyamot |
| **Date Created** | 2021-05-26, 10:35:27 AM |
| **Date Updated** | 2021-05-26, 11:16:21 AM |
| **Application** | Base Modification Analysis |
| **SMRT Link Version** | 10.1.0.119588 |
| **Inputs** | |
| **Path** | /net/eichler/vol28/projects/sequencing/pacbio/smrt-link/userdata/jobs_root/0000/0000007/0000007444 |

Analysis Parameters

##### Thumbnails

Mapped Polymerase Read Length

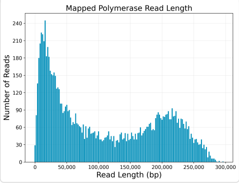


Coverage Across NC_013891.1.

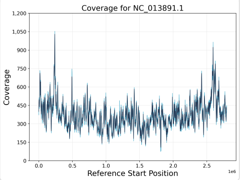


Kinetic Detections Histogram

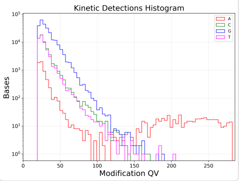


Alignment Length

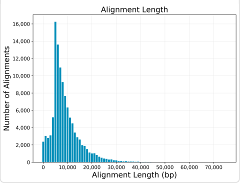


Depth of Coverage Distribution

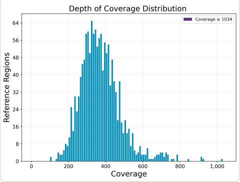


Modification QVs

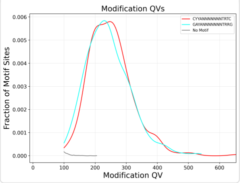


Alignment Concordance

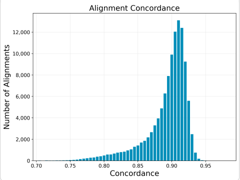


Coverage vs. [GC] Content

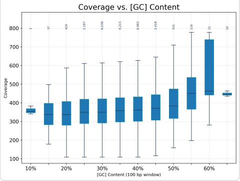


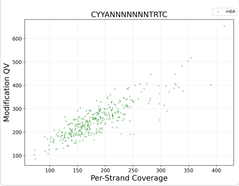


Mapped Concordance vs. Alignment Length

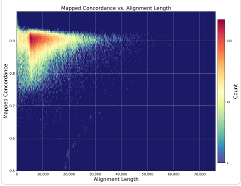


Per-Base Kinetic Detections

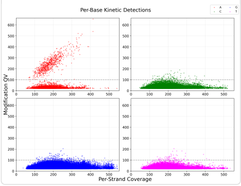


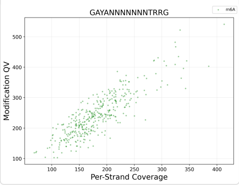


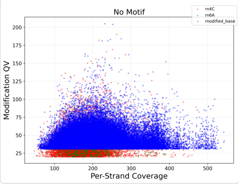

#### Mapping Report

| Value | Analysis Metric |
|---:|---|
| 96.63% | Percentage of Bases (mapped) |
| 108,100 | Number of Subreads (total) |
| 107,432 | Number of Subreads (mapped) |
| 668 | Number of Subreads (unmapped) |
| 99.38% | Percentage of Subreads (mapped) |
| 0.61% | Percentage of Subreads (unmapped) |
| 89.01% | Mean Concordance (mapped) |
| 1,062,527,186 | Number of Subread Bases (mapped) |
| 112,593 | Number of Alignments |
| 9,436 | Alignment Length Mean (mapped) |
| 11,108 | Alignment Length N50 (mapped) |
| 21,244 | Alignment Length 95% (mapped) |
| 75,916 | Alignment Length Max (mapped) |
| 9,945 | Number of Polymerase Reads (mapped) |
| 109,506 | Polymerase Read Length Mean (mapped) |
| 194,593 | Polymerase Read N50 (mapped) |
| 246,240 | Polymerase Read Length 95% (mapped) |
| 297,928 | Polymerase Read Length Max (mapped) |

#### Mapping Statistics Summary

| Sample | Movie | Number of Polymerase Reads (mapped) | Polymerase Read Length Mean (mapped) | Polymerase Read N50 (mapped) | Number of Subreads (mapped) | Number of Subread Bases (mapped) | Subread Length Mean (mapped) | Mea Con (ma |
|---|---|---|---|---|---|---|---|---|
| All Samples | All Movies | 9,945 | 109,506 | 194,593 | 107,432 | 1,062,527,186 | 9,436 | 89.0 |
| MC0513 MIX 383-5 | m64083_210514_071054 | 9,945 | 109,506 | 194,593 | 107,432 | 1,062,527,186 | 9,436 | 89.0 |

#### Mapped Polymerase Read Length

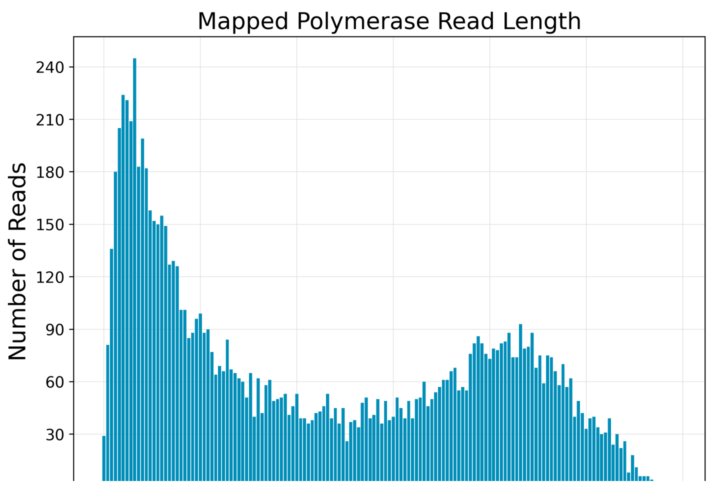

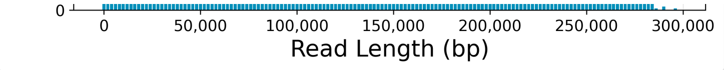


#### Alignment Length

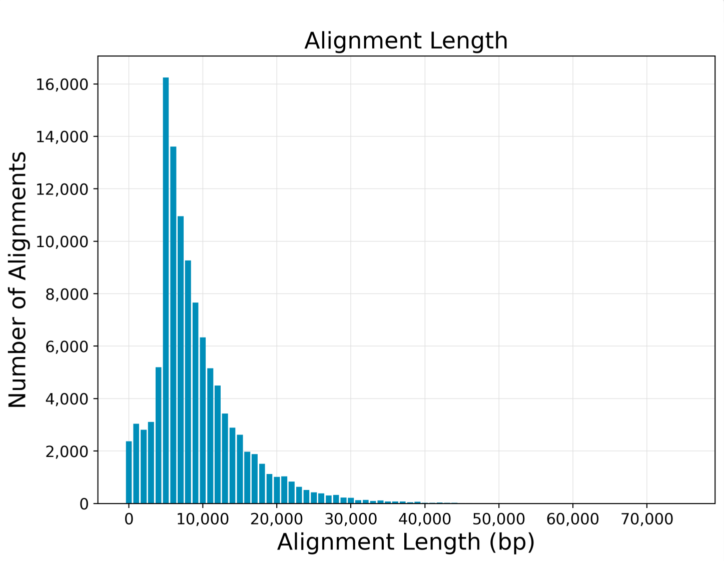


#### Alignment Concordance

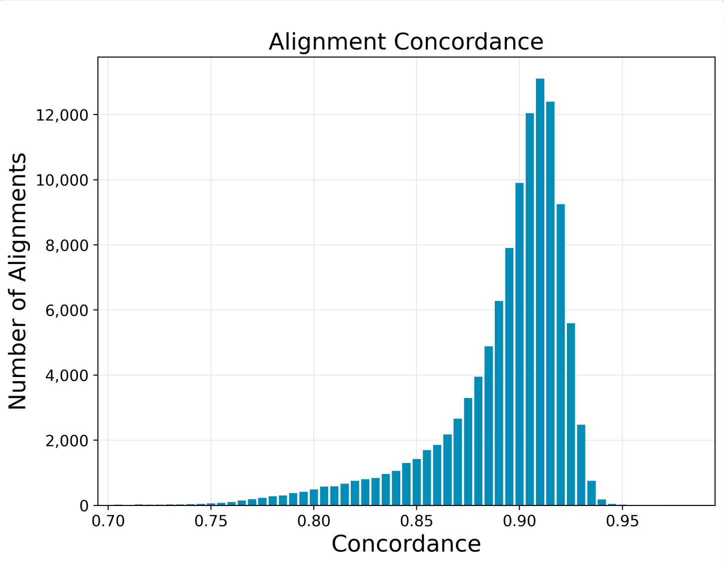


#### Mapped Concordance vs. Alignment Length

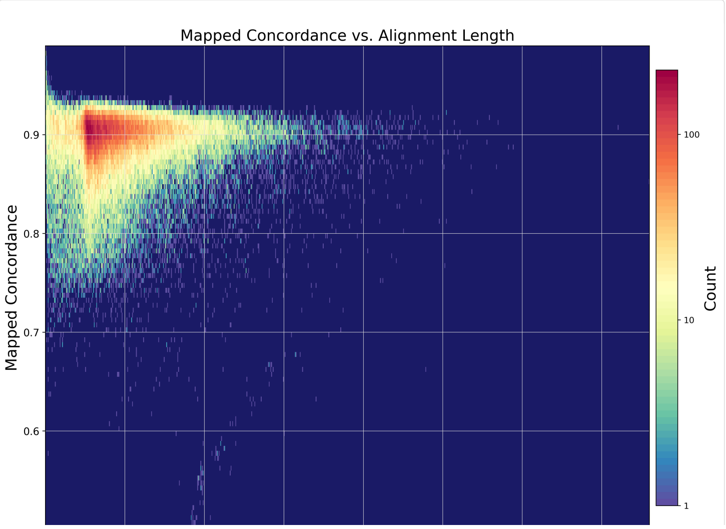

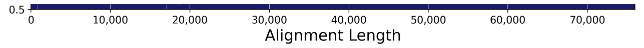


#### Coverage

| Value | Analysis Metric |
| --- | --- |
| 371 | Mean Coverage |
| 0.00% | Missing Bases |

#### Coverage Across Reference

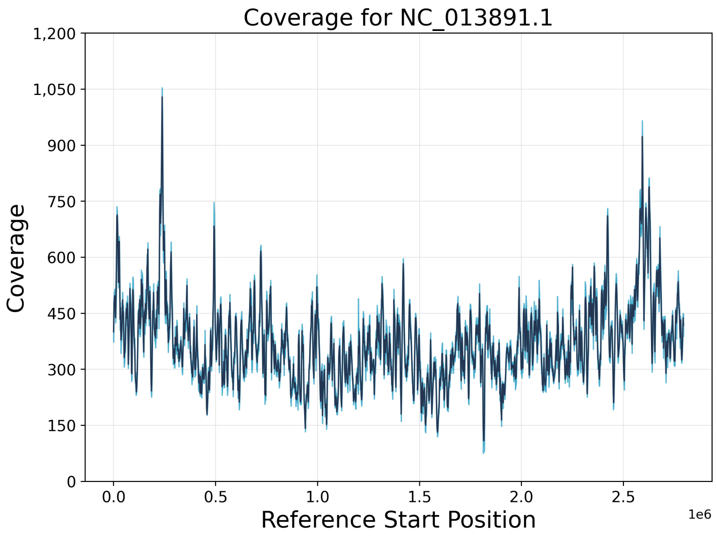


#### Depth of Coverage

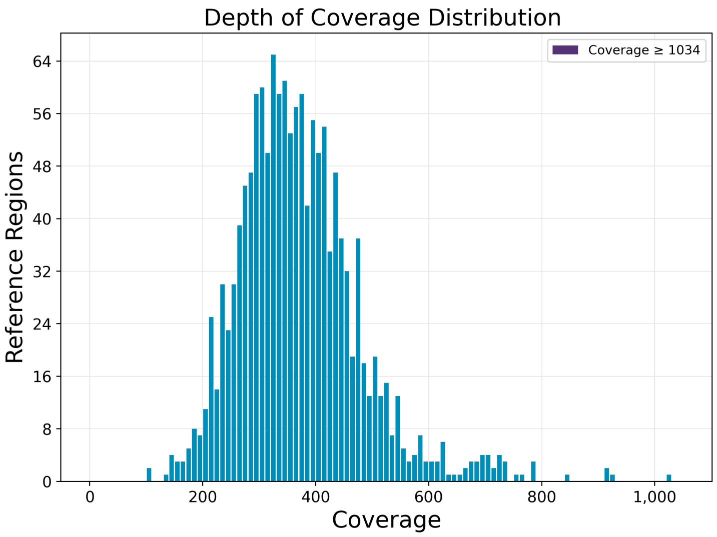


#### Coverage vs. [GC] Content

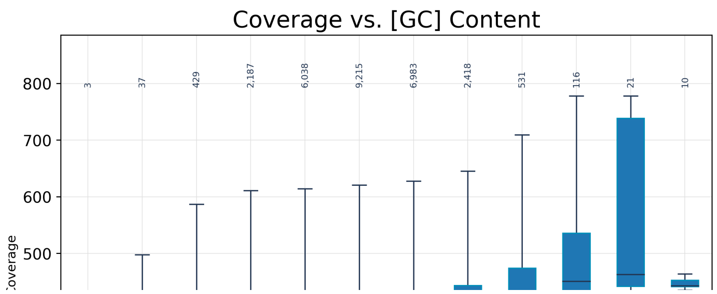

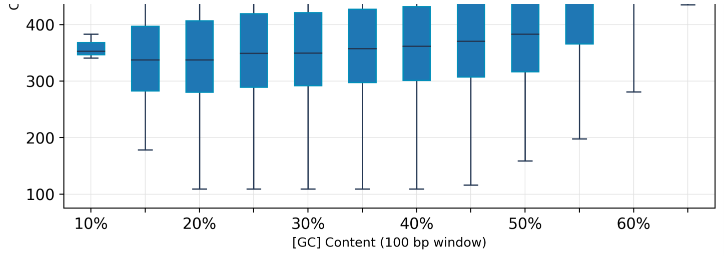


#### Kinetic Detections

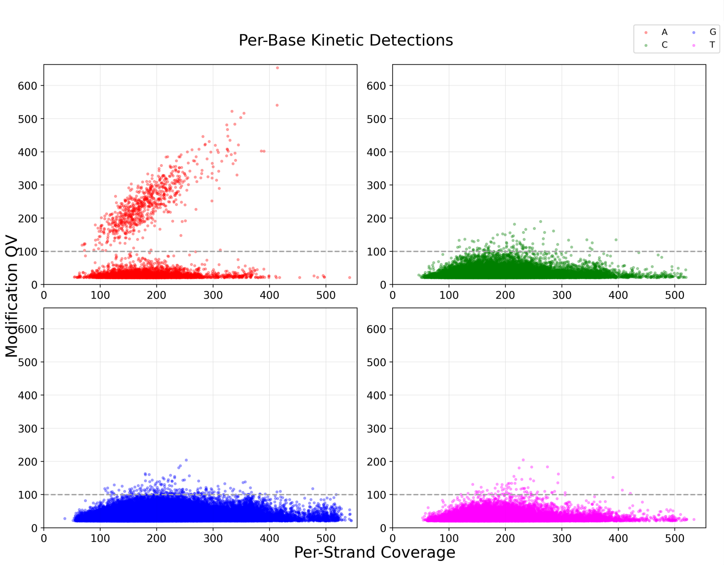


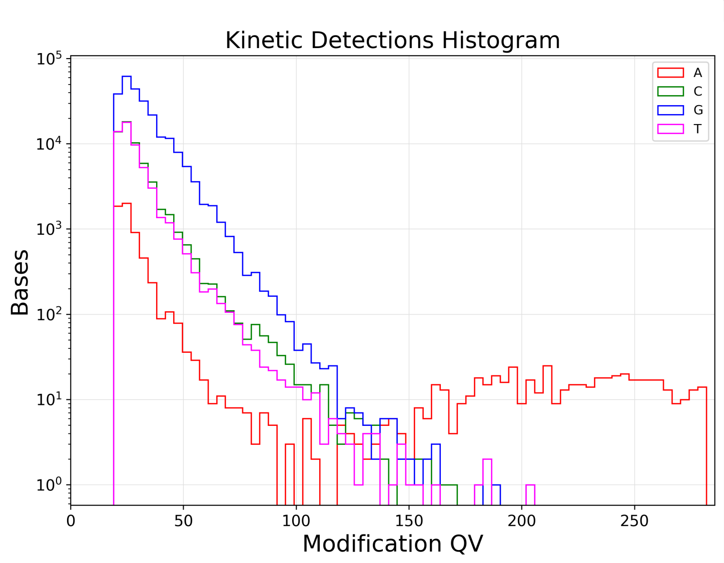


#### Modified Base Motifs

| Motif | Modified Position | Modification Type | % of Motifs Detected | # of Motifs Detected | # of Motifs in Genome | Mean QV | Mean Coverage | Partner Motif |
|---|---|---|---|---|---|---|---|---|
| CYYANNNNNNNTRTC | 4 | m6A | 99.4% | 362 | 364 | 251.2 | 183.9 | GAYANNNNNNNT |
| GAYANNNNNNNTRRG | 4 | m6A | 99.7% | 363 | 364 | 243.0 | 182.9 | CYYANNNNNNNTI |

#### Modification QVs

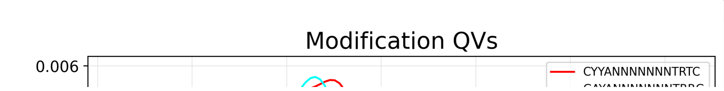

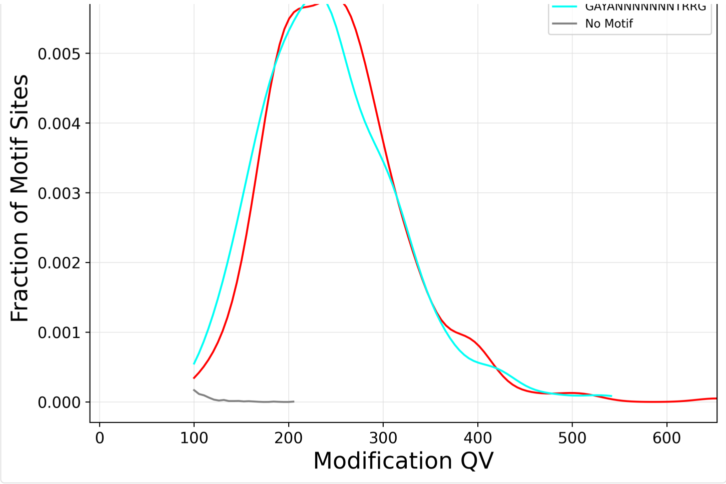


#### ModQV Versus Coverage By Motif

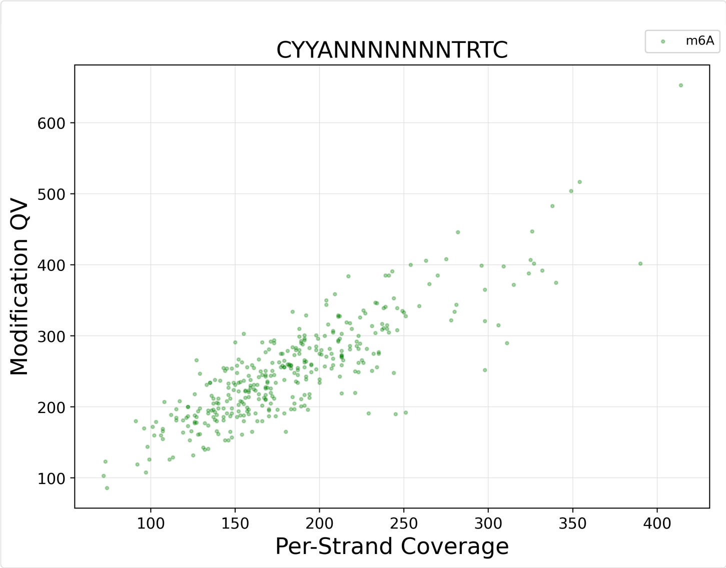


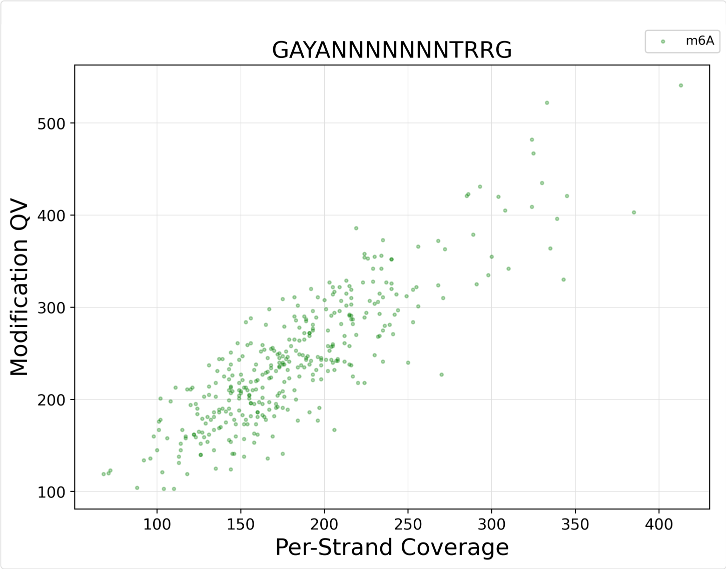


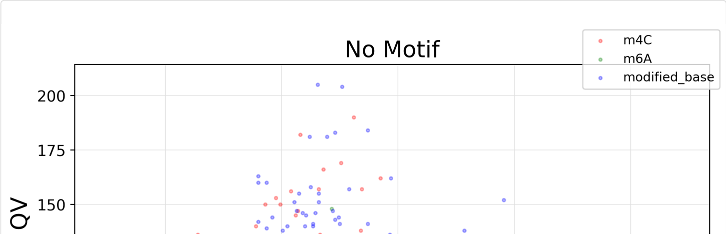

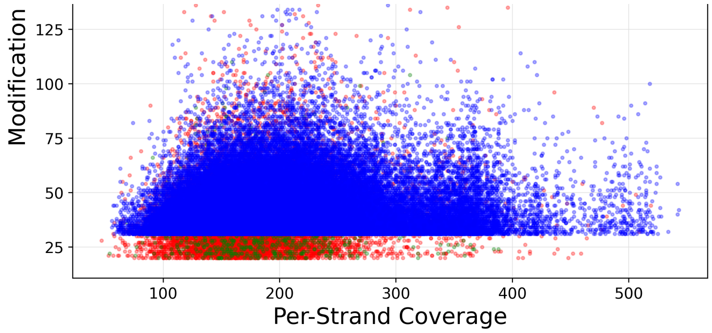


#### File Downloads

Edit Output File Name Prefix **Example:**analysis-DeltaRM2-7444

#### IGV Visualization Files

| File | Path | Size | Type |
|---|---|---|---|
| Mapped BAM Index | http://eee-smrt.gs.washington.edu:9090/job-data/pb_basemods/b7a9ea74-32f6-4db5-96dc-c7d7f8b7dbc3/call-auto_consolidate_alignments/execution/mapped.bam.bai | 192 KB | bam_bai |
| Mapped BAM | http://eee-smrt.gs.washington.edu:9090/job-data/pb_basemods/b7a9ea74-32f6-4db5-96dc-c7d7f8b7dbc3/call-auto_consolidate_alignments/execution/mapped.bam | 2 GB | bam |
| Per-Base IPDs for IGV | http://eee-smrt.gs.washington.edu:9090/job-data/pb_basemods/b7a9ea74-32f6-4db5-96dc-c7d7f8b7dbc3/call-gather_bigwig/execution/ipds.bw | 19 MB | bigwig |

###### SMRT Link Log

```
[INFO] [2021-05-26T10:35:32.471-07:00] Initial Setup of resources. Starting to run engine job id:7444 type:analysis Parent
MultiJobId:7442
[INFO] [2021-05-26T10:35:32.471-07:00] Job output directory is /net/eichler/vol28/projects/sequencing/pacbio/smrt-
link/userdata/jobs_root/0000/0000007/0000007444
[INFO] [2021-05-26T10:35:32.832-07:00] Final task options before converting to Cromwell inputs:
[INFO] [2021-05-26T10:35:32.835-07:00] [{
  "id": "run_find_motifs",
  "value": true,
  "optionTypeId": "boolean"
}, {
  "id": "consolidate_aligned_bam",
  "value": true,
  "optionTypeId": "boolean"
}, {
  "id": "dataset_filters",
  "value": "",
  "optionTypeId": "string"
}, {
  "id": "kineticstools_p_value",
  "value": 0.001,
  "optionTypeId": "float"
}, {
  "id": "kineticstools_compute_methyl_fraction",
  "value": true,
  "optionTypeId": "boolean"
}, {
  "id": "motif_min_score",
  "value": 100,
  "optionTypeId": "integer"
}, {
  "id": "motif_min_fraction",
  "value": 0.3,
  "optionTypeId": "float"
}, {
  "id": "mapping_min_concordance",
  "value": 70.0,
  "optionTypeId": "float"
}, {
  "id": "mapping_min_length",
  "value": 50,
  "optionTypeId": "integer"
}, {
  "id": "mapping_biosample_name",
  "value": "",
  "optionTypeId": "string"
}, {
  "id": "mapping_pbmm2_overrides",
  "value": "",
  "optionTypeId": "string"
}, {
  "id": "downsample_factor",
  "value": 0,
  "optionTypeId": "integer"
}, {
  "id": "kineticstools_identify_mods",
  "value": "m4C,m6A",
  "optionTypeId": "string"
}]
[INFO] [2021-05-26T10:35:32.836-07:00] Workflow options from job submission:
List(ServiceTaskIntOption(cromwell.workflow_options.max_nchunks,48,integer),
ServiceTaskStrOption(cromwell.engine_options.memory,32 G,string),
ServiceTaskIntOption(cromwell.workflow_options.nproc,8,integer),
ServiceTaskStrOption(cromwell.workflow_options.log_level,INFO,string),
ServiceTaskIntOption(cromwell.workflow_options.memory_per_core,4,integer),
ServiceTaskBooleanOption(cromwell.engine_options.read_from_cache,false,boolean),
ServiceTaskStrOption(cromwell.engine_options.backend,computecfg_00,string),
ServiceTaskBooleanOption(cromwell.engine_options.write_to_cache,true,boolean))
[INFO] [2021-05-26T10:35:32.836-07:00] Final engine options for analysis:
{"read_from_cache":false,"write_to_cache":true,"backend":"computecfg_00","default_runtime_attributes":
{"backend":"computecfg_00","maxRetries":1,"memory":"32 G","username":"kmiyamot"}}
[INFO] [2021-05-26T10:35:32.846-07:00] Submitting Cromwell workflow to http://localhost:9096
[INFO] [2021-05-26T10:35:32.910-07:00] Started Cromwell workflow b7a9ea74-32f6-4db5-96dc-c7d7f8b7dbc3 from SMRT Link job
5ede452a-427d-4440-ae79-bbd9d2fd88f8
[INFO] [2021-05-26T10:35:42.922-07:00] Cromwell workflow now in state SUBMITTED
[INFO] [2021-05-26T10:35:42.978-07:00] 0 tasks finished
[INFO] [2021-05-26T10:36:05.076-07:00] Cromwell workflow now in state RUNNING
[INFO] [2021-05-26T10:36:05.171-07:00] Created symlink to
/net/eichler/vol28/projects/sequencing/pacbio/nobackups/smrtlink_job_data/cromwell-executions/pb_basemods/b7a9ea74-32f6-4db5-
96dc-c7d7f8b7dbc3
[INFO] [2021-05-26T10:36:05.190-07:00] 1 task finished, task pb_basemods.update_subreads and 0 others running
[INFO] [2021-05-26T10:36:19.686-07:00] 3 tasks finished
[INFO] [2021-05-26T10:36:37.098-07:00] 5 tasks finished
[INFO] [2021-05-26T10:36:58.008-07:00] 6 tasks finished, task mapping.pbmm2_align and 4 others running
[INFO] [2021-05-26T10:38:29.013-07:00] 7 tasks finished, task mapping.pbmm2_align and 3 others running
[INFO] [2021-05-26T10:39:12.217-07:00] 9 tasks finished, task mapping.pbmm2_align and 1 other running
[INFO] [2021-05-26T10:40:03.979-07:00] 11 tasks finished, task mapping.gather_alignments and 0 others running
[INFO] [2021-05-26T10:41:04.275-07:00] 14 tasks finished, task coverage_reports.summarize_coverage and 2 others running
[INFO] [2021-05-26T10:42:04.630-07:00] 17 tasks finished, task pb_basemods.auto_consolidate_alignments and 22 others running
[INFO] [2021-05-26T10:44:05.332-07:00] 18 tasks finished, task pb_basemods.ipdsummary and 21 others running
[INFO] [2021-05-26T10:45:05.823-07:00] 21 tasks finished, task pb_basemods.ipdsummary and 18 others running
[INFO] [2021-05-26T10:46:06.475-07:00] 23 tasks finished, task pb_basemods.ipdsummary and 16 others running
[INFO] [2021-05-26T10:47:07.101-07:00] 29 tasks finished, task pb_basemods.ipdsummary and 10 others running
[INFO] [2021-05-26T10:48:07.585-07:00] 33 tasks finished, task pb_basemods.ipdsummary and 6 others running
```

```
[INFO] [2021-05-26T10:49:08.108-07:00] 39 tasks finished, task pb_basemods.ipdsummary and 0 others running
[INFO] [2021-05-26T10:50:10.803-07:00] 44 tasks finished, task pb_basemods.find_motifs and 1 other running
[INFO] [2021-05-26T10:52:11.644-07:00] 45 tasks finished, task pb_basemods.find_motifs and 0 others running
[INFO] [2021-05-26T11:14:18.663-07:00] 47 tasks finished, task pb_basemods.motifs_report and 0 others running
[INFO] [2021-05-26T11:16:19.375-07:00] Cromwell workflow now in state SUCCEEDED
[INFO] [2021-05-26T11:16:19.661-07:00] 48 tasks finished
[INFO] [2021-05-26T11:16:20.890-07:00] Cromwell workflow completed, importing 13 outputs
[INFO] [2021-05-26T11:16:20.927-07:00] about to write datastore for job 5ede452a-427d-4440-ae79-bbd9d2fd88f8
[INFO] [2021-05-26T11:16:20.970-07:00] Done writing datastore for job 5ede452a-427d-4440-ae79-bbd9d2fd88f8
[INFO] [2021-05-26T11:16:20.970-07:00] Successfully completed running core job id:5ede452a-427d-4440-ae79-bbd9d2fd88f8
[INFO] [2021-05-26T11:16:21.153-07:00] Imported 11 files
Linked 4 reports to 3 datasets
Job id:7444 DataStoreFile 3e150923-e1ad-849c-412a-1db44f6e342b imported
[INFO] [2021-05-26T11:16:21.169-07:00] Updated job 5ede452a-427d-4440-ae79-bbd9d2fd88f8 state to SUCCESSFUL
[INFO] [2021-05-26T11:16:21.195-07:00] Job 7444 type:analysis state:SUCCESSFUL is NOT eligible for email sending. no
createdByEmail user. Skipping sending Email
```
